# Dietary fibre responses emerge through host-dietary regime interplay: Apple pectin modulates immune reactivity in adult zebrafish

**DOI:** 10.64898/2026.10.29.762775

**Authors:** Justin Abi Assaf, Gwenaelle Le Gall, Ruth Sullivan, Anastasia Bezrucico-Pyatakov, Hanne Carlsson, Jayme Cohen-Krais, Jean-Charles de Coriolis, Alice May Godden, Daniel Marcu, Daniel Gorton, Simone Immler

**Affiliations:** School of Biological Sciences, University of East Anglia, Norwich Research Park, NR4 7TJ, Norwich, UK; Norwich Medical School, University of East Anglia, Norwich Research Park, NR4 7TJ, Norwich, UK

**Keywords:** Short-chain fatty acids, Liquid chromatography-tandem mass spectrometry, Lipopolysaccharide, Immune responses, Intestinal alkaline phosphatase

## Abstract

Dietary fibres are rarely consumed independently. Instead, they form part of an overall diet that includes dietary fibres from different sources, in varying amounts and consumed under different feeding patterns. Here, we tested how dietary regimes containing apple pectin, oat β-glucan, or cellulose shaped the host. We analysed gut microbiome composition by 16S rRNA sequencing, measured brain and intestinal SCFA profiles by targeted metabolomics, and assessed immunomodulation by measuring gene expression systemically in the blood and locally in the spleen, as well as intestinal enzyme activity. The dietary intervention was conducted for 12 weeks on adult female and male zebrafish (*Danio rerio*). Our results show that the gut microbial composition can be relatively stable across different dietary fibre exposure patterns yet still vary significantly between the sexes. Additionally, the effect of sex on the gut microbiome may become more evident when dietary fibres are administered at high dosages, but intermittently. Dietary fibre regimes can alter intestinal short-chain fatty acid profiles more readily than brain profiles, which show a stronger sexually dimorphic response. Interestingly, apple pectin modulated the zebrafish immune response through interactions dependent on the overall regime, tissue, sex, and host immune status. Notably, tissue *il1b* gene expression differed depending on how apple pectin was administered and the host’s physiological state. Collectively, our study emphasises the highly context-specific effects of dietary fibres, shaped by the dynamic interplay between the overall dietary regime and intertwined host factors.

**Graphical Abstract:** 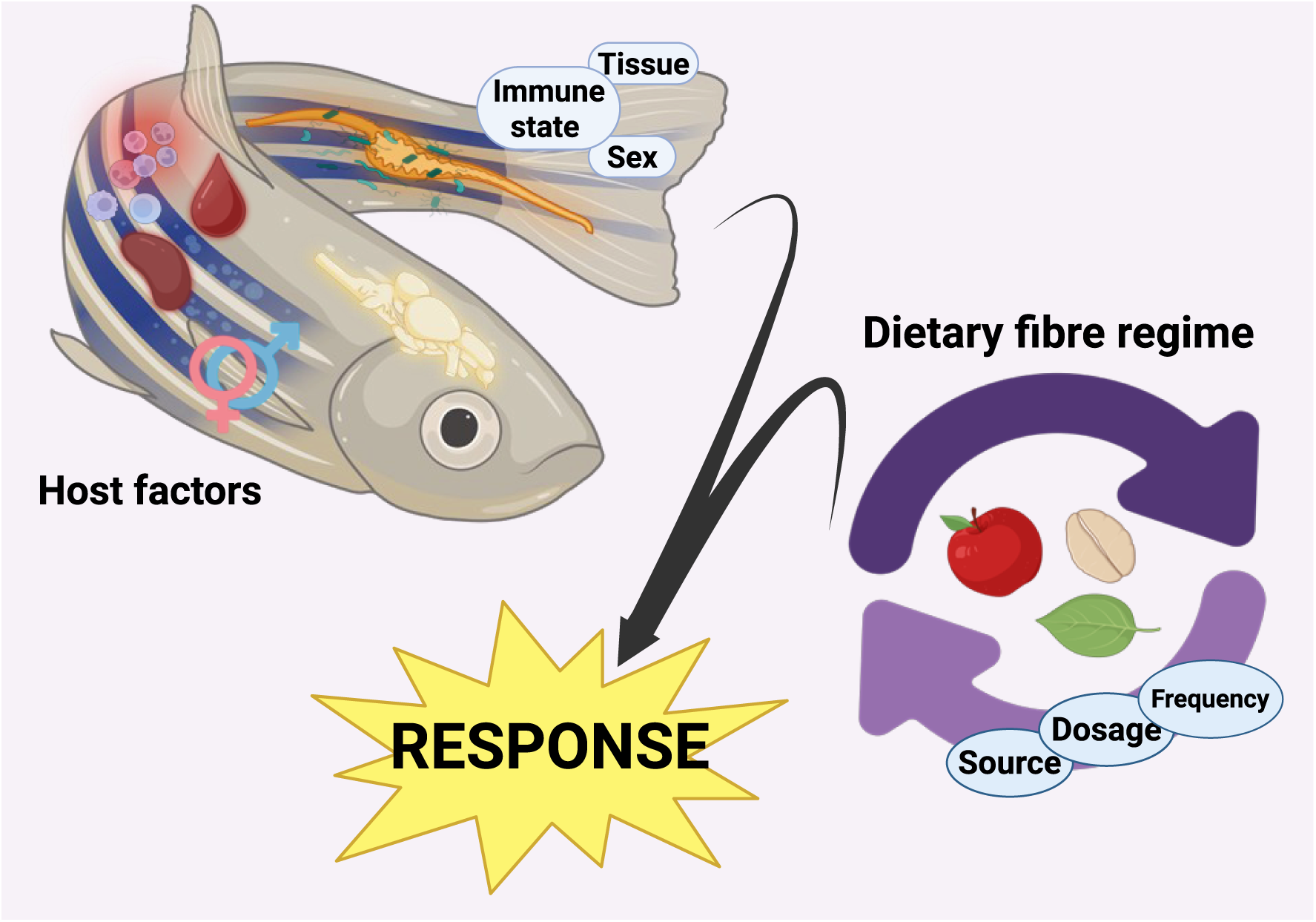

Created in https://BioRender.com

## INTRODUCTION

Dietary fibres (DFs) are a heterogeneous group of non-digestible polysaccharides whose bioactive properties are determined by their source, chemical structure, and physicochemical properties (solubility, viscosity, and fermentability)^1–3^. For this reason, the health effects of DFs can be difficult to disentangle, as they emerge from interactions between fibre type and dosage, the food matrix, feeding regime, intervention duration, host physiological state, and baseline gut microbiome^3–5^. They are also an essential component of a healthy diet and can modulate the gut-brain axis, metabolism, and the inflammatory response^6–8^. Although there is a movement towards plant-based diets rich in DFs, global guidelines for adult DF intake remain inconsistent. Some recommendations specify a blanket recommended daily amount, while others offer sex-specific guidelines^9^. Standardising DF intake is therefore challenging, as optimal requirements can vary significantly with factors such as age, sex, metabolism, and body composition^10^. The biological interplay between diet, nutrition, and the host is complex, resulting in context-dependent responses to DFs, which are conditional on the host’s physiological background or intrinsic state^5,11–13^. Females and males exhibit differences in gut microbiome structure, metabolism, and immune response^14,15^. Despite this, our understanding of the role and influence of sexual dimorphism in the incorporation of DFs into the diet is very limited. The literature suggests a sex-dependent response in mammalian models, including differences in microbial community restructuring^16^, fermentation and SCFA production^16,17^, and immune signalling^12,16^. However, the use of non-mammalian animal models, such as zebrafish (*Danio rerio*), in this context is underexplored.

In this study, we conducted a 12-week randomised block-design dietary intervention to test the effects of the dosage and frequency of three different DFs on fitness parameters in sexually mature female and male zebrafish. Purified, commercially available DFs were selected and incorporated into the basal Control diet. We selected two soluble, highly fermentable DFs, apple pectin (AP) and oat β-glucan (B), and one insoluble, minimally fermentable DF, cellulose (C). We specifically chose these DFs for their relevance to the human diet. For each of the three DFs, fish were subjected to either a 3% continuous low dosage (3 g fibre per 100 g basal diet, corresponding to a final inclusion level of 2.9% w/w in the complete diet) or a 15% intermittent high dosage (15 g fibre per 100 g basal diet, corresponding to a final inclusion level of 13% w/w in the complete diet). Specifically, we administered the 3% DF regime five times per week and the 15% regime once per week throughout the 12-week period. Rather than considering DF type and dosage in isolation, we also incorporated feeding pattern to better capture the broader dietary context in which DFs are consumed and to provide a temporal dimension. This allowed us to investigate whether the same DF, provided at a comparable overall exposure (12 weeks) but under different feeding regimes, could elicit distinct biological responses. Here, we aimed to determine whether the host response depends not only on what DF is consumed and how much, but also on how that exposure is distributed over time. In addition to the gut microbial and host metabolic responses to the various DF regimes, we also investigated their role in shaping and possibly supporting the immune system, both systemically (blood) and in localised immune compartments (spleen and intestine). After the dietary intervention, we measured blood and splenic gene expression of key immune response genes (*tnfa*, *il1b*, *il6*, and *myd88*) and quantified intestinal alkaline phosphatase (IAP) activity. For gene expression, we compared fish exposed to purified oral lipopolysaccharides (LPS), a potent immunostimulant^18^, mixed with the basal diet, with an LPS-negative cohort to assess the host response to the DF regimens across two distinct physiological states.

Here, we tested how the overall DF regime shaped the gut microbiome composition, gut and brain SCFA levels, and immune responses in female and male zebrafish. Using molecular, metabolic, and analytical approaches, we show that sexual dimorphism is a key driver of the physiological and genetic responses to DFs. We also show that the immunological response varies across tissues (blood, spleen, and intestine) depending on the dietary regime. Notably, our results also highlight that dietary regimens containing AP can modulate immune reactivity in a sex-specific manner across both genetic and enzymatic readouts. Altogether, our results support the idea that the responses to DFs are conditional and highly context-dependent, in part attributable to the host’s intrinsic biological status (e.g. sex, immune state, tissue). Additionally, the interplay between host and diet is shaped by the broader dietary context and the host’s physiological state, thereby contributing to variability in dietary responses. These findings pave the way for further studies exploring the dynamic interplay between host and diet, particularly as nutritional research increasingly shifts towards a more personalised approach.

## RESULTS

### Gut microbial community responses to dietary fibre regimes are influenced by sex

We housed the fish in mixed-sex tanks across the various dietary regimes, maintaining a 1:1 sex ratio per tank (**Fig. S1**). We assessed the impact of different DF regimes on gut microbial communities by 16S rRNA sequencing (LPS-negative cohort, n = 5 per sex and DF regime) to identify possible sex-specific shifts in our mixed-cohort tank setup.

Fusobacteriota was the dominant phylum across all dietary regimes and both sexes, accounting for over 53% of the taxonomic distribution, followed by Proteobacteria/ Pseudomonadota. At the genus level, *Cetobacterium* was the most abundant (based on relative abundance) genus of the phylum Fusobacteriota across all the regimes (between 53% and 73%) (**Fig. S2**). Our results show that the overall microbial composition was broadly consistent across the seven DF regimes; however, we found sex-specific amplicon sequence variants (ASVs) within each regime, suggesting that sex contributed more to microbial composition than the dietary regimes, even within the Control diet. The flower plot shows that, across all seven regimes, female and male fish shared only 38 core ASVs. Female fish under the Control, 15AP, 3C, and 15C regimes had more unique ASVs overall than their male counterparts. This pattern appears to be the inverse when comparing the sexes under 3AP, 3B, and 15B (**Fig. 1A**). We focused on the relative abundance of Fusobacteriota because it was the dominant phylum in our study. In the Control group, female fish had higher Fusobacteriota (69%) than males (63%). Both sexes under the 3% soluble fibre (AP and B) regimes showed similar Fusobacteriota relative abundance. However, under both 15AP and 15B, Fusobacteriota relative abundance was higher in males than in females. Also, Fusobacteriota was dominant in male fish compared to female fish under both cellulose regimes (3C and 15C) (**Fig. 1B** and **Fig. S3**). The data suggest a clear sex influence on microbiome composition and appear to shape the observed response to the dietary regime, especially in fish given the DF once a week at a higher dosage (i.e. under the 15% regime).

**Fig. 1:**
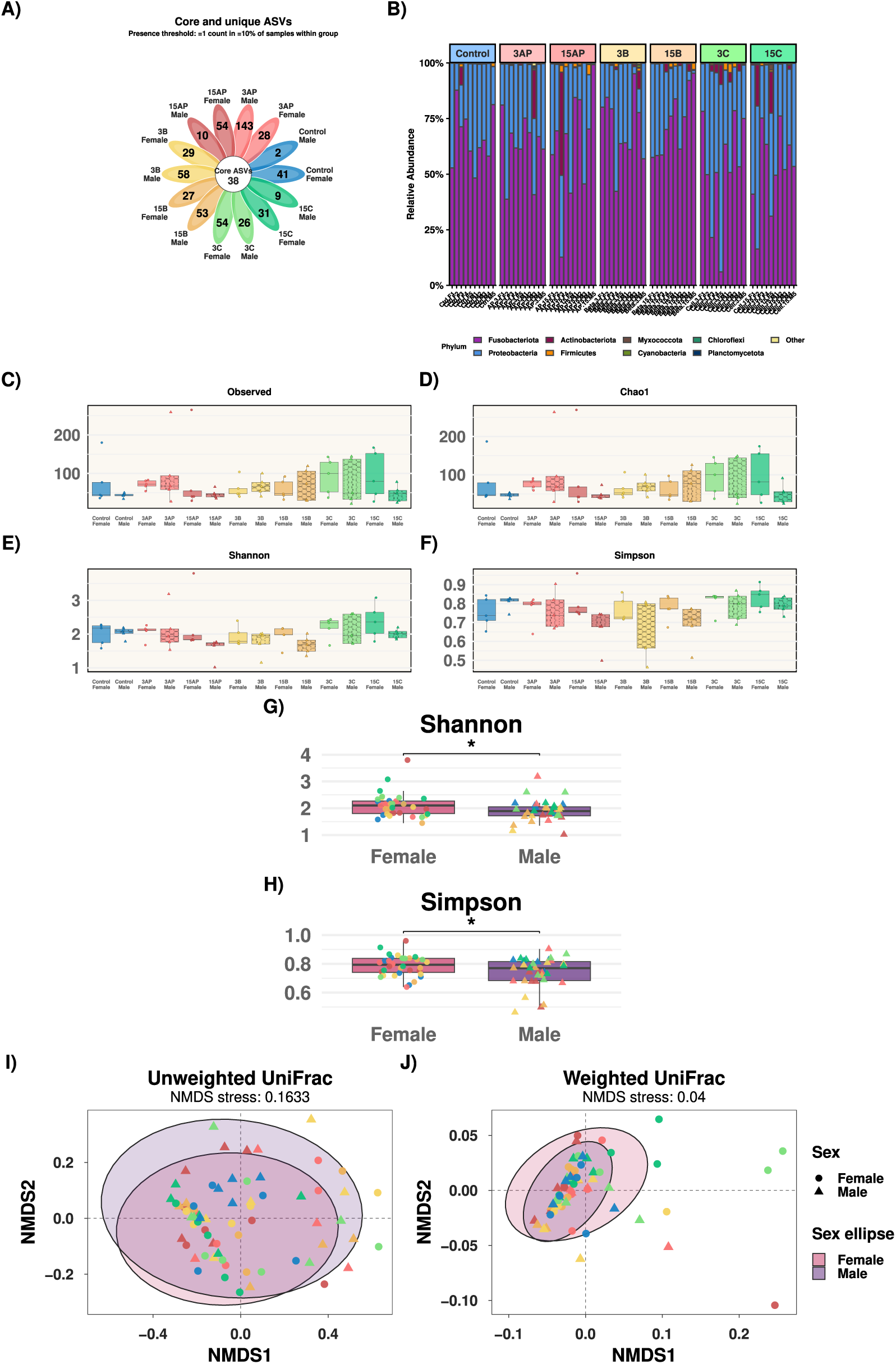
Gut microbial community responses to DF regimes stratified by sex. (**A**) Flower plot showing shared and unique ASVs. (**B**) Relative abundance plot. (**C-F**) α-diversity indices box plots. (**G-H**) Sex differences in α-diversity (Shannon and Simpson). (**I-J**) UniFrac phylogenetic β-diversity. Each point represents an individual sample. * denotes *p* < 0.05

After running a single-shot rarefaction (depth = 44,453), the DF regimes did not significantly affect the microbiome α-diversity (Kruskal-Wallis for “Diet”, *p* > 0.05) (**Fig. 1C-F**). To test for sex differences in the gut microbiome, we compared microbial diversity and richness using four α-diversity indices (Observed, Chao1, Shannon, and Simpson). For both Observed and Chao1 indices, the “Diet:Sex” interaction term was not statistically significant (lmer, *p* > 0.05), but the main effect of “Sex” was statistically significant for Shannon (*F*_1,37.943_ = 4.81, *p* = 0.034) and Simpson (*F*_1,38.98_ = 5.525, *p* = 0.024). Notably, female zebrafish exhibit significantly higher Shannon (estimated difference = 0.199 ± 0.091 SE, *p* = 0.034) and Simpson (estimated difference = 0.046 ± 0.02 SE, *p* = 0.024) α-diversity than male zebrafish, suggesting that females have a more diverse gut microbial community (a larger number of different taxa) as well as a more even distribution (less taxonomic dominance) than their male counterparts, irrespective of dietary regime and despite the mixed-cohort setup (**Fig. 1G and H**). We then explored β-diversity among samples across the various regimes using a phylogenetic approach. The unweighted UniFrac diversity (qualitative) was similar across “Diet”, “Sex”, and their interaction (PERMANOVA, 9999 permutations, *p* > 0.05). In contrast, weighted UniFrac diversity (quantitative) was statistically significant, driven by “Sex” (PERMANOVA, *F* = 3.74, df = 1, R^2^ = 0.049, *p* = 0.03). However, the main effect of “Diet” was marginally non-significant (PERMANOVA, *F* = 1.75, df = 6, R^2^ = 0.137, *p* = 0.07), suggesting a potential mild impact on β-diversity despite not reaching the statistical threshold (**Fig. 1I and J**). Next, we investigated whether feeding frequency modified the strength of the sex-associated microbiome pattern in the DF-supplemented fish (3AP, 15AP, 3B, 15B, 3C, 15C). We found that the sex-associated pattern is more evident in weekly fibre-treated fish (15AP, 15B, 15C) (PERMANOVA, *F* = 4.29, df = 1, R^2^ = 0.125, *p* = 0.01) compared to fish given the DFs five times a week (3AP, 3B, 3C) (PERMANOVA, *F* = 1.296, df = 1, R^2^ = 0.042, *p* = 0.286). This suggests that host-associated structure, such as sex, becomes more apparent when DFs are provided once a week rather than more frequently. However, we did not detect a statistically significant “Sex:Frequency” interaction (PERMANOVA, *F* = 0.64, df = 1, R^2^ = 0.01, *p* = 0.51); hence, the stratified results should not be viewed as confirmatory. To assess the impact of sex on weighted UniFrac diversity, we conducted a betadisper analysis, which quantifies the dispersion between the two sexes (the multivariate analogue of Levene’s test). We detected no significant difference in multivariate dispersion between females and males, supporting the interpretation that the weighted UniFrac sex effect was primarily driven by differences in microbial community rather than unequal within-sex variability (*F* = 2.13, df = 1, *p* = 0.15). To further determine if any specific taxa within each sex were responsible for driving the observed β-diversity difference, we conducted a differential abundance analysis using ANCOM-BC2 (Analysis of Compositions of Microbiomes with Bias Correction 2) at the genus level, with multiple testing using the Benjamini-Hochberg false discovery rate adjustment^19,20^. We detected no differentially abundant genera between experimental diets and Control, nor between the sexes (*q* > 0.05) (**Fig. S4**).

Altogether, the data show that across the seven dietary regimes, gut microbial communities remained broadly conserved, with sex explaining the dominant source of β-diversity variation. Sex-related differences were apparent at the community level rather than in specific taxa, potentially reflecting subtle variation distributed across multiple taxa.

### Dietary fibre effects on SCFA profiles were primarily intestinal, with brain profiles predominantly influenced by sex

SCFAs are major microbial fermentation products of DFs and key mediators of host physiology, with evidence that they can enter systemic circulation and interact with the blood-brain barrier (BBB)^21,22^. We quantified the microbial-derived SCFAs across the gut-brain axis to determine how SCFA profiles varied by dietary regimen and whether these responses were sex- and/or tissue-specific. Five linear SCFAs (acetate, propionate, butyrate, valerate, caproate) and four branched SCFAs (isobutyrate, 2-methylbutyrate, isovalerate, isocaproate) were detected in our samples. It is worth noting that 2-methylbutyrate, isovalerate, and isocaproate were poorly detected and had reduced analytical precision. We retained them in the dataset to provide a complete metabolic profile; however, their biological relevance should be interpreted with caution.

We tested whether different DF regimes affect microbial-derived SCFAs in the intestine, despite the strong sexual dimorphism in gut microbiome diversity observed in our previous analysis. The GI tract is the first organ to encounter digested food. We hypothesised that different dietary regimes would have a stronger effect on the intestinal SCFA profile in relation to host sex. We found that across all seven regimes and in both female and male intestines, acetate, valerate and propionate were the three most abundant intestinal SCFAs. In female intestines, acetate accounted for 85.6% of the total SCFA pool, valerate 6.4%, and propionate 5%. On the other hand, in male intestines, acetate accounted for 89.2% of the total SCFA pool, and propionate and valerate accounted for 5.27% and 2.74%, respectively.

Notably, “Diet” had a statistically significant effect on the three most abundant intestinal metabolites: acetate (*F*_6,21.49_ = 2.61, *p* = 0.047), valerate (*F*_6,83_ = 4.05, *p* = 0.001) and propionate (*F*_6,83_ = 2.65, *p* = 0.021) (refer to **Fig. S5**). Interestingly, fish under the 3AP regime had higher intestinal acetate, propionate, and valerate levels than the Control, the opposite of the other regimes. However, we detected no statistically significant change in intestinal acetate or propionate levels when comparing each experimental regime to the Control group (averaged over sex) (*p*_adj_ > 0.05). Only fish under the 15B regime had lower intestinal valerate compared to Control (*t*(83) = −2.69, *p*_adj_ = 0.044) (**Fig. 2A**). Additionally, we did not detect a significant sex effect on the three most abundant intestinal SCFAs, acetate, propionate, and valerate, except for marginally higher valerate in the female intestine than in the male intestine (*t*(83) = 1.967, *p* = 0.053).

**Fig. 2:**
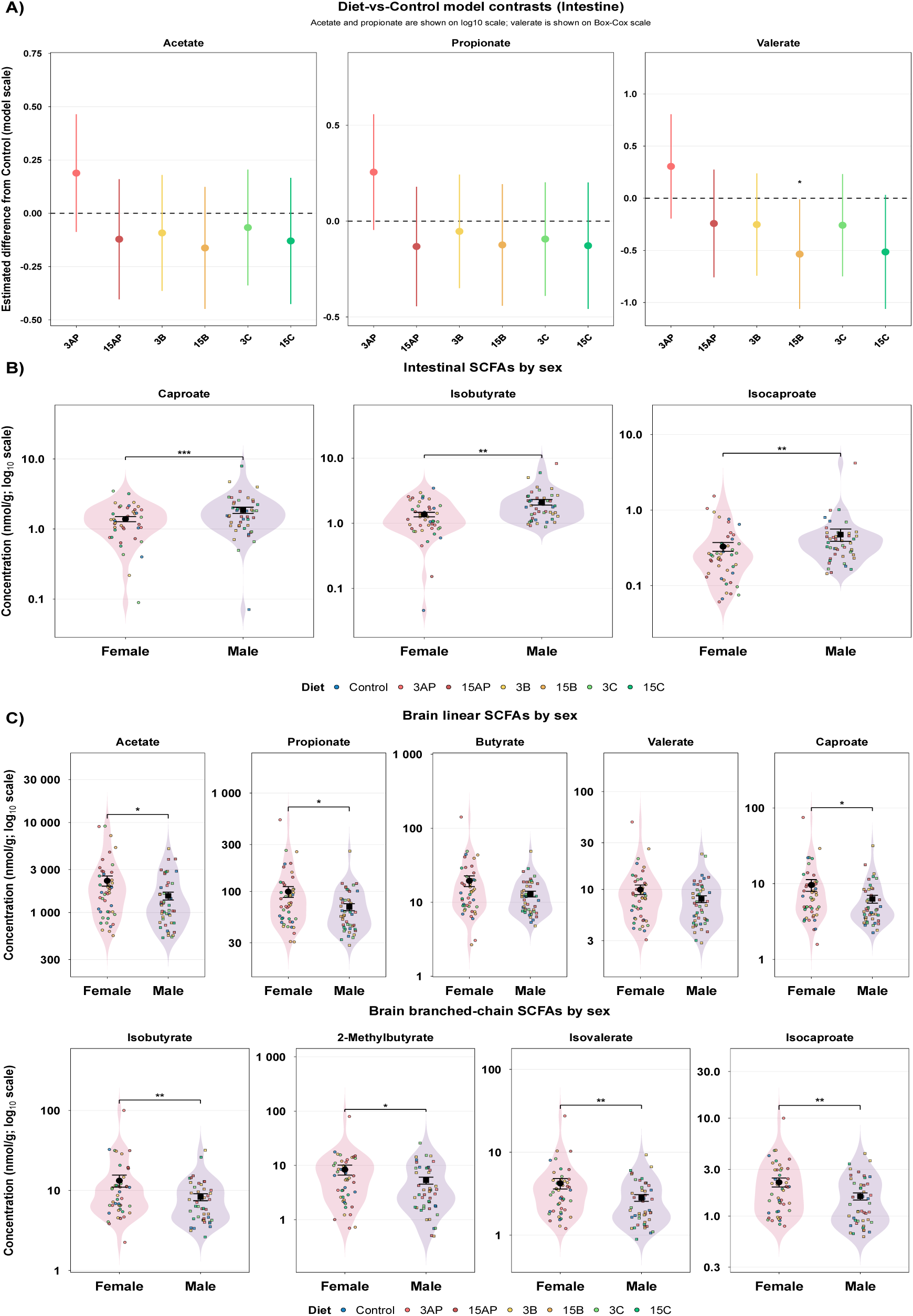
Intestine and brain SCFA concentrations. (**A**) Points represent model-estimated diet contrasts vs Control, with 95% confidence intervals. (**B-C**) Sex effects were estimated from the final model and averaged across diet levels. SCFA concentrations are displayed on a log10-transformed y-axis for better visualisation. Dark points represent group means, overlaid on individual sample points, and error bars represent ± SEM. For plotting simplicity, “Diet” refers to the dietary regime. * *p* < 0.05, ** *p* < 0.01.

In contrast, “Sex” significantly affected intestinal caproate levels (*F*_1,73.26_ = 12.17, *p* < 0.001), intestinal isobutyrate levels (*F*_1,82.69_ = 10.59, *p* = 0.002), and intestinal isocaproate levels (*F*_1,82.96_ = 9.7, *p* = 0.003). Averaged across all diets, the main model estimated that female intestines had 37% less caproate (nmol/g) (*t*(73.3) = −3.488, *p* < 0.001), 35.4% less isobutyrate (nmol/g) (*t*(82.7) = −3.25, *p* = 0.002), and 33% less isocaproate (nmol/g) (*t*(83) = −3.11, *p* = 0.003) compared to male intestines (**Fig. 2B**). Finally, we found that female intestines weigh significantly more than male intestines (n = 46 per sex; Welch *t*-test: *t*(70.08) = 6.92, *p* < 0.001) (**Fig. S7**).

In the brain, in both females and males and across all seven regimes, acetate, propionate, and butyrate were the three most abundant SCFAs, accounting for approximately 90%, 5%, and < 1% of the total SCFA pool, respectively. We followed a similar analysis workflow as for intestine and found that the effects of “Diet” and the “Diet:Sex” interaction on all nine SCFAs were not statistically significant (*p* > 0.05). However, most brain SCFAs differed significantly between females and males. “Sex” as a main effect was statistically significant on acetate brain levels (*F*_1,68.16_ = 5.4, *p* = 0.023), propionate brain levels (*F*_1,68.49_ = 6.27, *p* = 0.015), caproate brain levels (*F*_1,71.26_ = 5.15, *p* = 0.026), isobutyrate brain levels (*F*_1,69.97_ = 7.03, *p* = 0.010), 2-methylbutyrate brain levels (*F*_1,70.29_ = 5.60, *p* = 0.021), isovalerate brain levels (*F*_1,68.76_ = 7.07, *p* = 0.010), and isocaproate brain levels (*F*_1,68.5_ = 7.1, *p* = 0.010). However, brain butyrate and valerate did not differ by sex (*p* > 0.05) (refer to **Fig. S6**). We conducted a post hoc analysis, averaging across all seven dietary regimes to examine the distribution between females and males more specifically. Notably, the main model predicted that female brains contained 35% more acetate (nmol/g) than male brains (*t*(68.2) = 2.33, *p* = 0.02), 30% more propionate (nmol/g) than male brains (*t*(68.5) = 2.5, *p* = 0.01), 34% more caproate (nmol/g) than male brains (*t*(71.3) = 2.27, *p* = 0.03), 36% more isobutyrate (nmol/g) than male brains (*t*(70) = 2.65, *p* = 0.01), 47% more 2-methylbutyrate (nmol/g) than male brains (*t*(70.3) = 2.37, *p* = 0.02), 40% more isovalerate (nmol/g) than male brains (*t*(68.8) = 2.66, *p* = 0.01), and 35% more isocaproate (nmol/g) than male brains (*t*(68.5) = 2.66, *p* = 0.01) (**Fig. 2C**). On average, female brains weighed significantly less than male brains (n = 46 per sex; Welch *t*-test: *t*(87.52) = −2.36, *p* = 0.02) (**Fig. S7**).

Taken together, we find that the intestine is more metabolically responsive to DF regimes than the brain, which exhibits a more distinct sexually dimorphic metabolic profile. Additionally, sex-specific differences in both intestinal and brain SCFA profiles should be interpreted as an integrated host-microbe metabolic response, reflecting not only microbial activity but also host-dependent factors such as tissue physiology, metabolite uptake, utilisation, and systemic metabolism^23–25^.

### Blood immune responses to dietary fibre regimes are sex-specific

We investigated the effects of the DF regimes and sex on the expression of key immune response genes (*tnfa*, *il1b*, and *il6*) in the blood of female and male zebrafish. Blood provides a systemic readout of the host’s physiology and immune state, serving as a communication network between organs^26^. DFs can modulate immune gene expression via microbiota-dependent pathways, such as those involving SCFAs, and/or microbiota-independent interactions with host epithelial and immune cells^27,28^. Immune genes act sequentially as part of signalling cascade pathways, exhibiting a time-dependent and temporal expression pattern in which early mediators trigger downstream responses^29,30^. In our study, we assessed all gene expression levels 24 hrs after the fish ingested the dry feed with or without purified LPS.

We detected a statistically significant “Diet:Sex” interaction for all the three tested genes: *tnfa* (*F*_6, 251.22_ = 2.19, *p* = 0.04), *il1b* (*F*_6_,_255.17_ = 3.24, *p* = 0.004), and *il6* (General linear model: *F* = 4.92, df = 6, *p* < 0.001) (**Fig. S9**), emphasising that blood immune responses to DF is sex specific. Next, we conducted a post-hoc analysis comparing the gene expression between females and males within each of the seven DF regimes. Notably, females expressed lower *tnfa* than their male counterparts under the 3AP (log2FC = −1.82, *p* = 0.001), 15AP (log2FC = −1.14, *p* = 0.04), and 3C (log2FC = −1.272, *p* = 0.02) dietary regimes. The sex contrast for the other five regimes (Control, 3B, 15B, and 15C) did not meet the statistical significance threshold (*p* > 0.05) (**Fig. 3A**). Additionally, females expressed lower *il1b* levels compared to their male counterparts under the Control (log2FC = −1.052, *p* = 0.009), 3AP (log2FC = −0.866, *p* = 0.03), 15AP (log2FC = −0.962, *p* = 0.02), 15B (log2FC = −1.287, *p* = 0.002), and 3C (log2FC = −1.488, *p* < 0.001) (**Fig. 3C**). Similarly, females expressed lower *il6* levels than their male counterparts under the Control (log2FC = −1.374, *p* = 0.01), 3AP (log2FC = −1.736, *p* = 0.002), 15AP (log2FC = −1.304, *p* = 0.02), and 3C (log2FC = −1.45, *p* = 0.008). Interestingly, females under 15C had 3.7-fold higher *il6* expression than their male counterparts (log2FC = 1.895, *p* < 0.001) (**Fig. 3E**). This reversal suggests that the relationship between sex and *il6* expression depends on the overall dietary context of cellulose. We then conducted another post hoc analysis comparing each experimental regime with its same-sex Control group. We did not detect a statistically significant difference in expression relative to the Control for either sex (*p* > 0.05), except that *il6* expression was higher in 15C females compared with Control females (log2FC = 1.863, *p*_adj_ = 0.004). Interestingly, this response was evident under intermittent high-dosage cellulose (15C), but not under the 3C regime (**Fig. 3B, D, and F**).

**Fig. 3:**
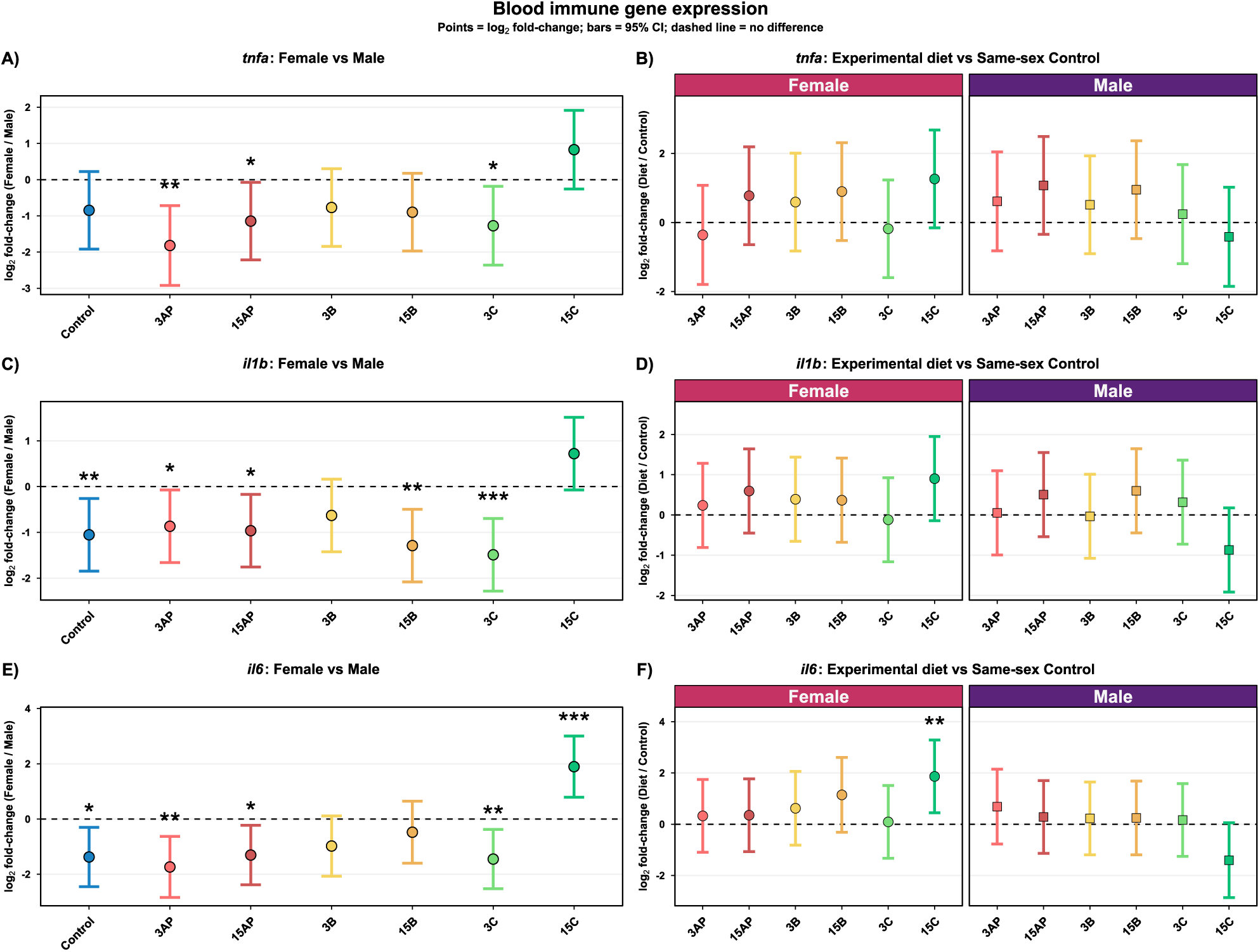
Blood *tnfa*, *il1b*, and *il6* expression by qPCR. (**A, C, and E**) Female vs male contrast for the specified gene. (**B, D, and F**) Experiment diet vs Control contrast, stratified by sex. Each point is the model-estimated log2FC, and the vertical error bar is its 95% confidence interval. * *p* < 0.05, ** *p* < 0.01, and *** *p* < 0.001. For plotting simplicity, “Diet” refers to the dietary regime.

Together, our results indicate that DF strongly modulates blood immune gene expression by sex, with adult female zebrafish typically exhibiting lower gene expression than their male counterparts. We also highlight that DF effects on blood gene expression can be linked to both composition and frequency.

### Dietary fibres shape the blood transcriptional response to LPS

Intraperitoneal or intrarectal LPS injection in adult zebrafish has previously been shown to induce an inflammatory response^31–33^. To investigate a potential interplay between DFs and LPS on the immune response, we analysed gene expression at the end of the dietary intervention, comparing fish orally administered LPS to an LPS-negative control group. We found that the effect of LPS was not statistically significant on *tnfa* expression (*F*_1,253.99_ = 0.031, *p* = 0.86), *il1b* expression (*F*_1,257.84_ = 0.004, *p* = 0.95), or *il6* expression (*F* = 0.053, df = 1, *p* = 0.82). However, oral LPS administration appeared to modulate blood *il1b* gene expression, with variation by dietary regime (*F*_6,256.13_ = 2.87, *p* = 0.01). Specifically, *il1b* gene expression was higher in 15AP fish given oral LPS than in their LPS-negative counterpart (log2FC = 1.373, *p* < 0.001) (**Fig. 4A**). Moreover, 15AP fish (LPS-positive) showed significantly higher *il1b* expression than the Control fish (LPS-positive) (log2FC = 1.405, *p*_adj_ = 0.003) (**Fig. 4B**). These observed changes in blood *il1b* appear unique to the 15AP regime and LPS, as this interplay does not affect the blood gene expression of the other cytokines, *tnfa* and *il6* (**Fig. S10**).

**Fig. 4:**
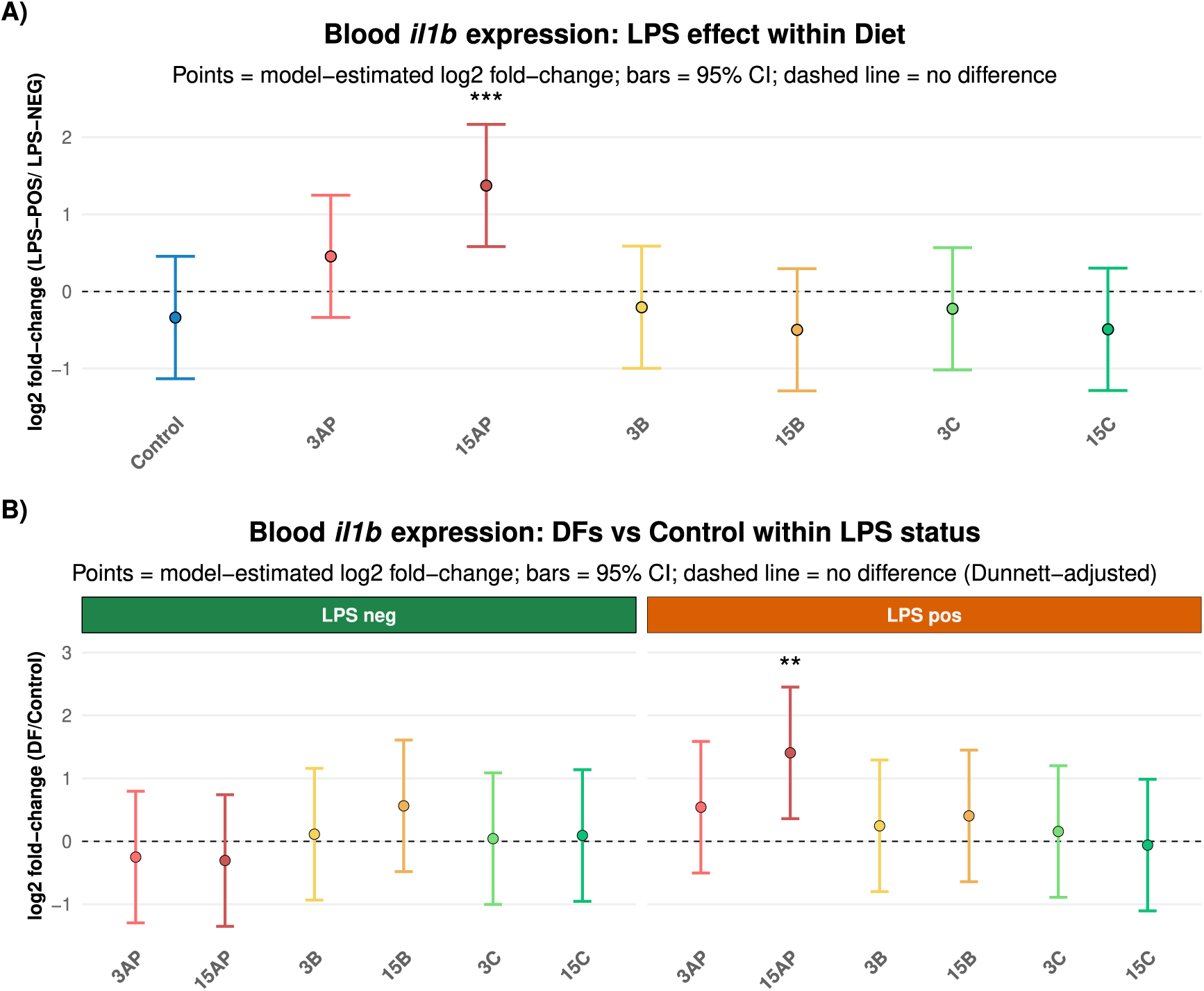
Blood *il1b* expression in response to LPS across dietary regimes. (**A**) Blood *il1b* expression comparing LPS-positive to LPS-negative within each diet. (**B**) Blood *il1b* expression comparing experimental to Control, stratified by LPS status. Each point is the model-estimated log2FC, and the vertical error bar is its 95% confidence interval. * *p* < 0.05 and ** *p* < 0.01. For plotting simplicity, “Diet” refers to the dietary regime.

### Apple pectin modulates immune reactivity in a sex- and tissue-specific manner

To further investigate how DFs shape the immune response, we then assessed the expression of *tnfa*, *il1b*, *myd88*, and *il6* in the zebrafish spleen, a major secondary lymphoid organ involved in both innate and adaptive immune responses^34,35^. Based on the previous blood gene expression results indicating an interplay between AP and LPS, we then conducted a follow-up qPCR experiment on fish from the Control, 3AP, and 15AP regimes across both LPS cohorts.

As in the blood, we detected a significant “Sex” main effect for splenic *tnfa* expression (*F*_1,97.834_ = 6.66, *p* = 0.01). Specifically, female spleen *tnfa* expression was lower than that of males (log2FC = −0.71, *p* = 0.01), averaged by “Diet” and “LPS” ( **Fig. S11**). We found no significant two-way interactions between “Diet” and “Sex”, “Diet” and “LPS”, or “LPS” and “Sex” for splenic *tnfa* expression (*p* > 0.05).

Interestingly, for splenic *il1b* expression, there was a significant three-way “Diet:LPS:Sex” interaction (*F*_2,100.2_ = 3.31, *p* = 0.04), indicating that the relationship between “Diet” and “LPS” on spleen *il1b* expression differed between females and males. To dissect this interaction, we conducted three post hoc analyses across three strata. (1) First, we tested the main effect of “Diet” within Sex:LPS combinations. Apple pectin-supplemented females and males (3AP and 15AP) from either LPS cohort showed different spleen *il1b* expression than their respective Control groups. However, in all cases, statistical significance was not met (*p*_adj_ > 0.05) (**Fig. 5A**). (2) Next, we explored the main effect of “LPS” within the Diet:Sex interaction. The oral administration of LPS did not induce a significant change in spleen *il1b* expression compared to the LPS-negative cohort in any of the Diet:Sex combinations (*p*_adj_ > 0.05) (**Fig. 5B**). (3) Finally, for each dietary regimen (3AP or 15AP), we examined whether its effect in comparison to Control varied with LPS status across the two sexes (interaction contrast). We found that neither 3AP nor 15AP significantly altered the Diet-vs-Control effect on spleen *il1b* following LPS exposure in either sex (*p*_adj_ > 0.05) (**Fig. 5C**). Overall, the results indicate that spleen *il1b* expression varies with the interaction among the DF regimen, LPS status, and sex. In contrast, the “Diet:LPS” interaction on blood *il1b* expression did not differ by sex.

**Fig. 5:**
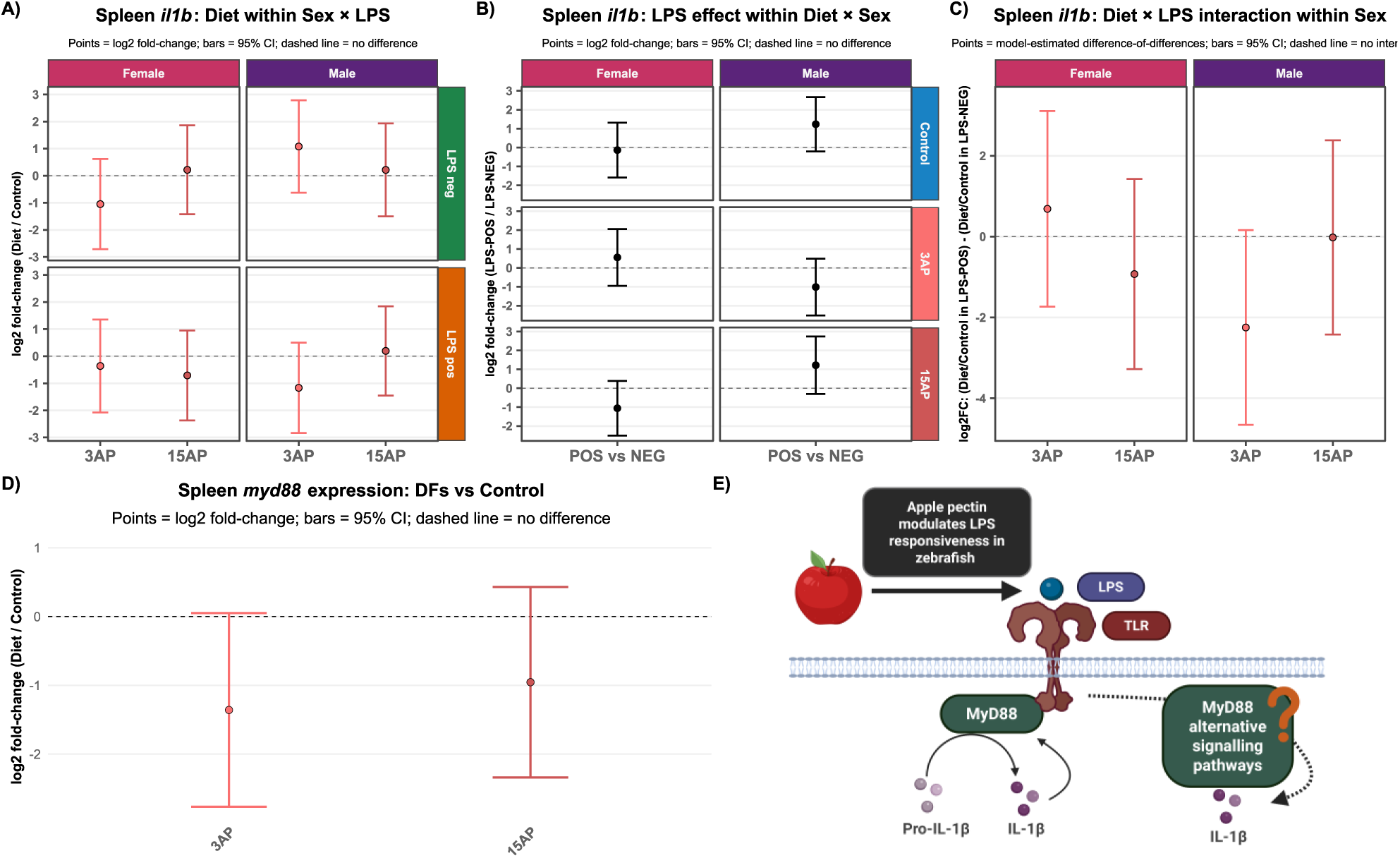
Spleen *il1b* and *myd88* expression by qPCR. (**A-C**) Stratified contrasts of spleen *il1b* expression dissecting the Diet × LPS × Sex interaction. (**D**) Spleen *myd88* expression comparing experimental to Control. (**E**) Proposed schematic involving the regulation of *il1b* by MyD88 in response to LPS. Each point is the model-estimated log2FC, and the vertical error bar is its 95% confidence interval. For plotting simplicity, “Diet” refers to the dietary regime.

In the follow-up experiment, we tested spleen *myd88* gene expression as an additional proxy for innate immune signalling, as it contributes to downstream inflammatory responses, including pathways associated with *il1b* induction in response to stimuli such as LPS^36–38^. We detected a marginally significant “Diet” effect on the expression of *myd88* (*F*_2,16.336_ = 2.98, *p* = 0.08). No significant effects of “Sex” or “LPS”, or any two-way interactions, were detected (*p* > 0.05). In the post hoc analysis, when comparing both apple pectin regimes (3AP and 15AP) to the Control, fish expressed lower spleen *myd88* than the Control. However, the differences did not reach statistical significance (*p*_adj_ > 0.05) (**Fig. 5D**). The findings suggest that the interplay between AP and LPS on spleen *il1b* expression, previously detected, could involve alternative pathways beyond changes in MyD88-associated signalling (non-canonical pathways)^39–42^ (**Fig. 5E**).

Lastly, spleen *il6* gene expression remained unchanged when testing the main effects of “Diet”, “Sex”, “LPS” and their interactions across all conditions (*p* > 0.05) (**Fig. S11**). Since we administered LPS orally, we investigated diet-associated localised immunomodulatory effects in the intestine by quantifying intestinal alkaline phosphatase (IAP) activity using a fluorometric-based assay. Here, we tested only fish from the LPS-positive cohort, given the bidirectional, synergistic relationship between IAP and LPS (gut microbiome-derived or exogenous), whereby LPS can stimulate IAP activity as a defence mechanism, and IAP can neutralise excessive LPS, thereby maintaining homeostasis^43–45^.

First, we screened all seven dietary regimes and found that only the main effect of “Sex” was statistically significant for IAP activity (*F*_1,39_ = 24.875, *p* < 0.001). Notably, the post hoc analysis showed that IAP activity was significantly lower in female fish than in male fish, averaged across all seven regimes (*t*(39) = −4.99, *p* < 0.001) (**Fig. 6A**). Additionally, IAP under each experimental dietary regime did not differ statistically from their same-sex Control (*p*_adj_ > 0.05) (**Fig. 6B**). Overall, our results suggest that dietary regimes including AP are the most immunomodulatory. Limited evidence from a study in rats suggests that pectin can increase or restore IAP activity, particularly under diet-induced obesity^46^. Thus, we next focused on quantifying IAP activity in fish in Control, 3AP, and 15AP regimes. We observed a statistically significant “Diet:Sex” interaction (*F*_2,64_ = 3.18, *p* = 0.048). Given the borderline significance (α = 0.05), we further tested this interaction and conducted additional post hoc tests. (1) First, we compared IAP activity in the experimental groups (3AP or 15AP) with their same sex Control but found no statistical significance (*p*_adj_ > 0.05) (**Fig. 6C**). This suggests that the main effect of the dietary regimes on IAP was not strong enough to induce significant changes, suggesting a potential sex influence in response. (2) We then explored whether the IAP activity between the experimental regimes (3AP and 15AP) and the Control group differed between females and males. Indeed, we found that the sex difference in dietary response (3AP vs Control) was larger in females than in males (*t*(64) = 2.4, *p* = 0.02) (**Fig. 6D**). The data emphasise that the magnitude of the 3AP regime’s effect on IAP activity is sex-dependent, with female fish showing a stronger response than male fish. We did not detect a difference in dietary response (15 AP vs Control) between the sexes (**Fig. 6D**).

**Fig. 6:**
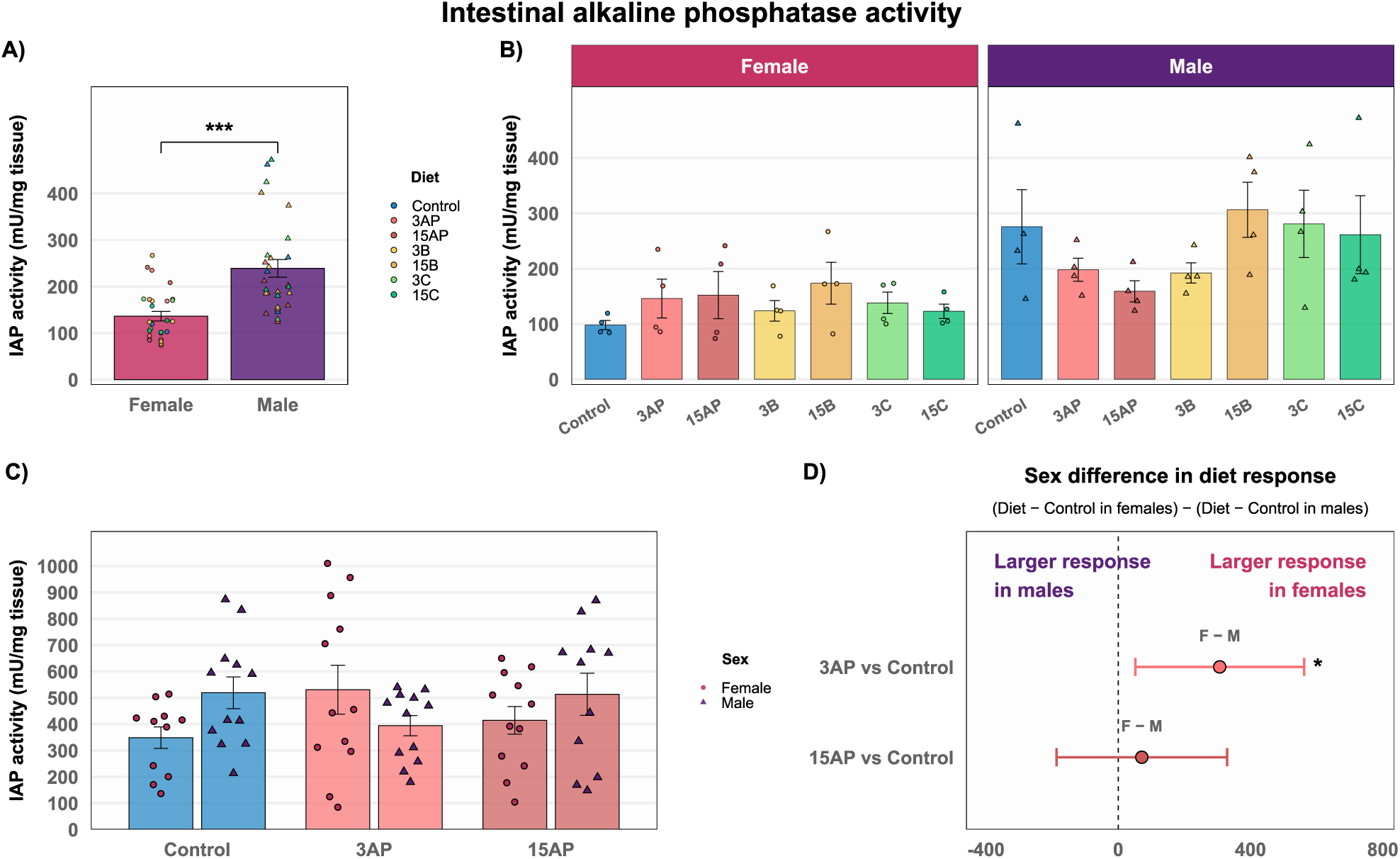
Intestinal alkaline phosphatase activity. (**A**) IAP activity (mU/mg tissue) by sex. (**B**) The initial screening of IAP across all seven regimes and stratified by sex. (**C**) IAP activity in the Control, 3AP, and 15AP regimes. (**D**) Model-estimated Diet × Sex interaction; points represent estimated responses and vertical error bars indicate 95% confidence intervals. * *p* < 0.05 and *** *p* < 0.001.

Overall, our findings indicate that adding AP to the zebrafish diet can influence immune responses at both genetic and enzymatic levels, though these effects vary greatly by context. In particular, the immune response to LPS is compartmentalised differently across tissues and is sex specific. Furthermore, AP’s ability to modulate the immune system depends on the overall dietary regime and the host’s immune status.

## DISCUSSION

In recent years, there has been growing interest in personalised nutrition to provide tailored dietary recommendations based on the host’s genotype, metabolic phenotype, health status and gut microbiome composition^47–50^. This framework reflects a broader recognition that dietary responses are context dependent. The physiological effect of a nutrient or dietary component is shaped not only by its intrinsic properties but also by the host’s physiological status and gut microbial context^51,52^. Here, we demonstrate that the effects of DFs are not confined solely to microbiome-mediated effects but are also shaped by the interplay between the dietary regime and sexually dimorphic characteristics in host physiology, tissue metabolism, and immune status. Collectively, these findings support a context-dependent view of DF function rather than a uniform one, highlighting that their health effects arise from complex host-diet-gut microbiome interactions.

We did not detect a statistically significant shift in gut microbiome composition due to DFs in any dietary regime, even in the fermentable ones (AP and B). Despite “Diet” having a larger effect size in weighted UniFrac β-diversity, “Sex” was the statistically significant predictor of microbial community changes. Achieving a statistically significant effect on the gut microbiome under controlled conditions depends heavily on the characteristics of the dietary intervention, including diet composition and matrix, dosage, feeding kinetics, and intervention duration^53^. Evidently, sex can shape the response to dietary interventions across vertebrate species, including humans, with effects extending beyond changes in gut microbial composition to influence metabolic and physiological outcomes^15,54,55^. Yet, consistent baseline differences in gut microbiome composition between female and male zebrafish have not been firmly established, with studies reporting variable sex-associated patterns depending on experimental conditions^25,56^. Here, we show that sex shapes the physiological context in which DF-associated shifts in the microbial community emerge, despite the sexes sharing the same tank environment and, theoretically, a shared microbiome composition through co-housing and faecal material^57,58^. Interestingly, the effect of sex on microbiome composition appears to be more pronounced when DFs were provided intermittently and at a high dosage. This suggests that the influence of host sex on the microbiome becomes highly apparent when dietary signals are subtle, which is aligned with research demonstrating that DF-induced changes are often more readily detectable at the gut microbiome level when fibre exposure is sustained and sufficient^59–62^. More broadly, the response to DFs is shaped not only by the characteristics of the dietary intervention itself, but also by the host-associated factors and the overall host-diet-gut microbiome interactions^63^. Thus, the contribution of diet and host to gut microbiome composition is dynamic, rather than static.

Compared with humans and mammalian animal models, data on SCFAs in zebrafish are scarce, especially in the context of DF interventions. In mammals, microbial-derived SCFAs and other metabolites act both locally in the gut and systemically in peripheral organs to regulate epithelial function, cellular proliferation, metabolism, and immunity^64–67^. However, despite putative evidence of conserved mechanisms and effects^68–71^, we still know little about their tissue distribution and kinetics, particularly in adult female and male zebrafish. We found that the intestinal and brain SCFA profiles did not show the same trend, highlighting tissue-specific SCFA regulation and compartmentalisation. Our results suggest that SCFAs are dynamic and that their contribution to gut-brain communication involves host physiology, gut microbiome composition, intestinal uptake rates, and systemic circulation. Metabolomic quantification of SCFAs represents a snapshot profile of the tissue and does not capture metabolic kinetics, including cellular uptake and utilisation^72,73^. Nevertheless, we show that the effects of the different DF regimes on SCFAs are more pronounced in the intestine than in the brain. Notably, we found that the 3AP regime was the only treatment to show a consistent numerical increase in intestinal acetate, propionate, and valerate compared to Control. However, these differences did not reach statistical significance (**Fig. 2B**). Interestingly, this suggests that soluble and fermentable DFs like AP should be consumed more consistently, rather than less, to potentially sustain intestinal SCFAs production^74,75^. Moreover, the data indicate that a DF’s capacity to generate SCFAs is not determined solely by fermentability. Instead, microbial fermentation depends on the overall physicochemical properties of a DF and the food source, both of which are important for generating SCFAs^76,77^. Pectin and cereal β-glucans are both soluble, fermentable dietary fibres that the gut microbiome can metabolise to generate SCFAs^77,78^. Accordingly, AP and B can both act as microbial substrates eliciting distinct fermentation dynamics, SCFA profiles, and host responses. Instead, neither 3B nor 15B displayed a similar pattern to 3AP on intestinal acetate, propionate, and valerate. Our data emphasise the complexity of DFs and intestinal SCFA production and show that, for a specific response to occur, several interrelated conditions involving the host, the overall diet, and the gut microbiome must be met. We also show the prevalent sexually dimorphic SCFA profile in the zebrafish brain. In line with the literature, in both mammals and zebrafish, the brain is a key somatic organ that exhibits sex-specific characteristics across molecular, hormonal, metabolic, neuroendocrine, and behavioural levels^79–82^. Our data indicate that the brain shows a more pronounced sexually dimorphic metabolic response than the intestine, which is more amenable to dietary changes. Unlike intestinal SCFAs, data on the mammalian brain SCFA profile, especially sex differences in concentrations, are difficult to obtain. We have previously shown using nuclear magnetic resonance (NMR), that acetate levels become particularly elevated in zebrafish brains rather than the intestine after an intervention with gum Arabic (GA)^79^. In this study, we employed targeted metabolomics using liquid chromatography-tandem mass spectrometry (LC-MS/MS). We found that the majority of quantified brain SCFAs, including acetate, were more abundant in female zebrafish than in their male counterparts across all dietary regimes (**Fig. 2D**). We cannot yet extrapolate these observations to humans, given the field’s infancy. However, the results open interesting avenues in gut-brain axis research and merit further study of the microbiome-dependent and independent effects of DFs on the brain.

The longstanding view in nutrigenomics and nutritional immunology is that adequate nutrient intake can support the immune system, whereas inadequate intake can compromise defence mechanisms^83–85^. Bioactive dietary components such as vitamin D^86–88^, polyphenolic compounds^89,90^, and omega-3 fatty acids^91–93^ have been shown to modulate immune cell function and induce epigenetic and transcriptomic changes, potentially leading to a more regulated and resolved immune response. In this study, we investigated the effects of DFs across different regimes on shaping the immune response, particularly in key immune organs, in both sexes and under two immune states: physiological baseline state (LPS-negative) and an LPS-exposed state (LPS-positive). Unlike female mammals, which typically exhibit stronger immune responsiveness than males^94–98^, we show that zebrafish immune gene expression, particularly in the blood, is higher in males than in females (**Fig. 3A, C, and D**). This aligns with immunology research in teleost fish, suggesting that the immune response is less directionally consistent than in mammals and is context-dependent (species, stimuli, and tissue)^99–101^. In aquaculture, LPS is typically used to activate the immune system, thereby improving defence and survival against future infections, similar to the use of vaccines^102,103^. Here, we administered a relatively high oral dosage of LPS to zebrafish to investigate whether this could prime the immune system or even induce acute inflammation. Interestingly, after LPS administration, we did not detect any signs of inflammation in the blood or spleen, at least at the genetic level (**Fig. S9** and **Fig. S11**). This observation suggests that when LPS is administered enterically, it primarily engages the gut mucosal immune system in adult zebrafish^43,104,105^, thereby possibly reducing its systemic dissemination and effects.

A key finding from our study was that adding AP to the zebrafish diet (3AP or 15AP regime) influenced *il1b* gene expression in the blood and spleen in response to LPS, as well as IAP activity. We highlight a distinct interaction between AP and LPS that shapes the zebrafish immune tone, suggesting that AP may prime subsequent responsiveness to LPS, with this response depending on the dietary regime, sex, and tissue. Pectin and related polysaccharides modulate the immune response, altering the expression or secretion of pro-inflammatory cytokines such as TNF-α, IL-1β, and IL-6 in a context-dependent manner^106–109^. Circulating immune cells offer a rapid, dynamic readout of systemic immune activation via fluctuating gene expression^110^. Our results suggest that when AP is administered intermittently at a high dosage (15AP) to zebrafish, blood immune cells will exhibit a greater cytokine response (specifically *il1b* expression) than naive immune cells that have not been previously exposed to AP (Control group) (**Fig. 4A and B**). We speculate that the 15AP regime may have induced a primed innate immune state in the blood, which became apparent only following LPS challenge. The *il1b* is an early marker of innate immune activation^111^, and its transcriptional response in zebrafish is detected early post-stimulus, often peaking within 3-6 hrs post-stimulation^111,112^. Also, *il1b* transcription in zebrafish usually declines by 24 hrs following a transient stimulus, unless there is active bacterial stimulation^113–115^. Interestingly, *il1b* expression remained detectable in both blood and spleen 24 h after LPS administration, suggesting a prolonged or enhanced inflammatory response. This state of immune priming could potentially confer protective effects against future pathogenic infections; yet, experimental validation is needed. Unlike the blood, where immune cells dynamically encounter circulating antigens, the spleen serves as a localised site for immune surveillance, filtering blood-borne antigens and encountering components and microbe-derived molecules that may originate in the intestine^34,116^. Our findings indicate a varied immune response in the spleen compared with the blood, emphasising how immune responses differ between tissues. We found that when AP is administered more frequently at a lower dose (3AP), male splenic *il1b* gene expression is lower in 3AP vs Control after LPS administration than in the absence of LPS, although this difference did not reach statistical significance (**Fig. 5A**). Our data show that the immune response is compartmentalised in a tissue- and sex-specific way and indicate that AP (depending on the regimen) modulates the zebrafish immune response. AP can either prime the immune system (evident by blood *il1b* expression) (**Fig. 4**) or act as an immune checkpoint, possibly reducing pathological hyperresponsiveness to a stimulus such as LPS (spleen *il1b* expression) (**Fig. 5**). Moreover, in rodents and zebrafish, pectin has been shown to increase IAP gene expression and/or activity^46,117,118^, and in zebrafish, IAP has been shown to detoxify the effects of LPS^43,44^. In this study, we did not detect a statistically significant change in IAP under the AP regimes (3AP and 15AP) compared to Control. Instead, we show that female zebrafish respond more strongly than males under the 3AP regime for IAP activity (**Fig. 6D**). It should be noted that in this study, we used a high degree of esterification AP (50-75%). High-degree esterification pectin has previously been shown to have a more context-dependent or dual immunomodulatory profile compared to low-esterified pectin, which exhibits a more explicit anti-inflammatory effect^119–124^. Moreover, unlike AP, regimes containing oat β-glucan (3B and 15B) showed no clear evidence of immunomodulatory activity under the tested conditions. This is consistent with research indicating that fungal and yeast β-glucans exhibit stronger immunomodulatory effects than cereal β-glucans^125^. Overall, these results emphasise that understanding the health impacts of DFs depends on analysing the complex interactions between their inherent physicochemical characteristics, dietary context, and the host’s physiological environment.

## CONCLUSION

In conclusion, this study shows why the biological response to DFs is difficult to determine and cannot be easily established. The overall DF regime can exert distinct downstream effects on the host depending on the sex, tissue, and immune context. Yet, host factors also shape the context through which DF effects arise. We show that sex shapes gut microbiome composition, with host-related effects becoming more evident under less frequent DF exposure, at least at the gut microbiome level. Despite no clear differences in tissue SCFA concentrations compared with the Control group, the effects of DFs on SCFAs varied by tissue, with diet-related effects apparent in the intestine, whereas sex-related differences were more evident in the brain. Finally, we show that apple pectin supplementation can modulate the zebrafish immune responses. However, these responses vary by the overall DF regime, tissue, immune context, and sex. Thus, DF effects depend on the dynamic interplay between dietary context and the host, with specific ramifications appearing only under certain conditions and remaining undetectable under others. Ultimately, establishing a clear causal effect of DFs on health requires a more complete understanding of the full range of possible interactions, and the conditions under which these interactions arise.

## METHODS

### Zebrafish husbandry and formulation of diets

ABWT zebrafish were obtained from the Zebrafish International Resource Center (ZIRC) at the University of Oregon, USA. The experimental fish used in this study were adult fish (6-8 months) reared from fertilised eggs obtained from crosses of F0 parents. The fish were housed in 3.5 L tanks (8 fish/tank), with images of gravel underneath each tank, in a recirculating rack system (ZebTEC rack with Active Blue technology, Tecniplast, UK), and kept on a 14:10 hour light:dark cycle. Water temperature was maintained at 28 °C with a pH of 6.5-8.5 and conductivity between 300 and 800 μS. The tank adult baffles were washed, and any excessive algae were removed every 3-4 weeks during the dietary intervention period. Health monitoring was performed daily during feeding. Experiments were performed with approval from the UK Home Office under project licence: PP1967239.

Six experimental diets were prepared by supplementing the standard dry feed/Control diet (Zebrafeed 400-600 by SPAROS, Área Empresarial de Marim, Lote C, 8700-221 Olhão, Portugal) with the different DFs at either 3% (2.9% w/w) or 15% (13% w/w). The same batch of Control diet (23.FO.CE.2411.ZF) was used throughout this study (**Table S1)**. The fish were fed at 2.5% of their body weight. For eight fish per tank, the DFs were added to the same amount of Control diet, as follows: (0.1 g Control feed + 0.003 g DF) for the 3% regime and (0.1 g Control feed + 0.015 g DF) for the 15% regime. The pectin from apple (product number 93854) (50 - 75% degree of esterification) and oat β-glucan (medium viscosity, product number P-BGOM) were obtained from Sigma-Aldrich (St. Louis, MO, USA) and Megazyme (Bray, Ireland), respectively. The insoluble dietary fibre cellulose (product number C6288) was also obtained from Sigma-Aldrich. Briefly, the DFs were supplemented into the Control diet by first weighing the desired amount, then adding a small amount of distilled water to create a slurry. This mixture was homogenised and then spread thinly before drying in an oven (50 °C) until bone dry. The next day, the dried sheets were placed in the freezer for a couple of days, then ground in a coffee grinder. The particles were sieved through a 400 μm mesh (Fisher Scientific, Loughborough, Leicestershire, UK). The prepared ground diets were stored in airtight containers in the freezer (−20 °C) until use.

### Experimental design

After acclimatisation, a total of n = 448 healthy adult fish (derived from multiple clutches) were randomly assigned to 56 3.5 L tanks and maintained at a 1:1 sex ratio (4 females and 4 males, n = 8 per tank). Each tank was randomly assigned one of the DF regimes and an LPS status (negative or positive). The experiment was divided into four blocks, with each block consisting of 14 tanks (one Control diet tank and 6 DF regimen tanks per LPS status). We used a full factorial design, combining each DF regime with LPS-negative and LPS-positive conditions. The DF regimes comprised Control; 3% and 15% apple pectin (AP); 3% and 15% β-glucan (B); and 3% and 15% cellulose (C). We measured the weight of eight fish per tank before and after the dietary intervention.

In the morning, the experimental fish were fed the supplemented dry feed either five times a week (Monday to Friday) on the low-dosage/habitual-consumption regime or once a week (Wednesday) on the high-dosage/intermittent-consumption regime. In the afternoon, live *Artemia franciscana,* from the Great Salt Lake, Utah, USA (Sep-Art Artemia Cysts, Ocean Nutrition), was provided daily for enrichment. Control fish were fed the standard dry feed seven times a week, in addition to *Artemia* in the afternoon. All the fish were fed 2.5% of their body weight. The weekday dry feed was standardised by pre-weighing and then storing in labelled Eppendorf tubes as follows: 0.1 ± 0.01 g for eight fish (Control), 0.103 ± 0.001 g for eight fish (low dosage/habitual consumption) and 0.115 ± 0.001 g for eight fish (high dosage/intermittent consumption). All fish were fed the prepared Control and experimental diets (rather than standard husbandry feed during the weekends) at 2.5% of their body weight throughout the study. The tank water flow was turned off for 10 min to allow undisturbed feeding before switching it back on. Each experimental block lasted for 12 weeks (**Fig. S1**).

### Acute oral administration of LPS

Lipopolysaccharide (LPS) from *Pseudomonas aeruginosa* (serotype 10.22, ATCC 27316), purified by phenol extraction, was purchased from Sigma-Aldrich. At the end of the 12-week dietary intervention, before the dissection day, the fish were either acutely exposed to LPS-supplemented feed or to regular feed without LPS (within 24 hrs before dissection). These feeds were administered twice daily at 6% of the zebrafish body weight (an adult zebrafish weighs approximately 0.5 g), with a 4-hr gap between the first and second feed, and without live *Artemia*. Feeding occurred pairwise based on their assigned dietary condition. The LPS-supplemented feed was prepared in sterile water at a concentration of 250 µg LPS/g fish per feed. Zebrafeed 400-600 was first weighed to account for the eight fish per tank and then mixed with the prepared LPS stock (5 mg/mL) in a large Petri dish. The food was then allowed to air-dry in a biosafety cabinet for approximately 4 hours. Then, the food was ground manually using a mortar and pestle before being coated with a few drops of menhaden fish oil (F8020-500ML) purchased from Sigma. After proper and careful mixing to ensure even coating, the food was stored at 4-6 °C in the fridge until use. The LPS-negative feed was prepared similarly but mixed with sterile water instead. For each block, the food was freshly prepared each week.

### Intestinal alkaline phosphatase (IAP) assay

For the initial intestinal alkaline phosphatase (IAP) activity screen, 56 intestinal tissues from the LPS-positive cohort were selected (n = 4 per sex per regime). The tissues were first weighed using an analytical scale (Fisherbrand), and the alkaline phosphatase fluorometric assay kit (product number: ab83371) (Abcam, Cambridge, UK) was used according to the manufacturer’s instructions, running two technical replicates for each sample. While on ice, the intestinal tissues were briefly washed with cold PBS, then homogenised with a pestle and centrifuged at 4 °C to collect the supernatant. After diluting the supernatant (1:100), they were incubated for 30 min in the dark at room temperature with the 4-methylumbelliferyl phosphate disodium salt (MUP) substrate in black 96-well plates for fluorescence-based assays. Then, the reactions were quenched with the “Stop Solution”. Finally, the Relative Fluorescence Units (RFU) were measured (excitation/emission = 360/440 nm) using the VANTAstar microplate reader (BMG LABTECH, Offenburg, Germany). IAP activity for each sample was derived from a standard curve (accounting for the dilution factor) and then normalised to its respective weight (mg). To minimise plate effects, the samples were randomly allocated to plates, maintaining an even balance of sex and diet. For the targeted analysis (Control, 3AP, and 15AP) (n = 12 per sex per regime), IAP activity was quantified in two independent runs, and the mean of the two runs for each sample was used in the analysis.

### Quantification of brain and intestinal SCFAs using LC-MS/MS

Around 0.75 g of 1.44 mm zirconium beads (Fisherbrand) were added to each of the 1.5 mL conical microtubes with screw-on O-ring cap (Fisherbrand). The beads and the tubes were left to chill on dry ice. The zebrafish intestinal and brain tissues (n = 7 per sex per diet per tissue) from the LPS-negative cohort (refer to **Fig. S8**), stored at −80 °C, were thawed on ice before weighing on an analytical scale (Fisherbrand). Then, the individual tissues were placed inside the pre-chilled tubes containing the beads. For each diet, intestinal and brain tissues were obtained from the same individual fish, ensuring that both organs reflected the same dietary exposure and reducing the intra-variability noise. Additionally, the selected samples per diet were chosen whilst maintaining an even Block distribution as much as possible. Next, 500 µL of 80% ice-cold methanol (80% MeOH + 20% molecular-grade water) was added to each tube. Samples were homogenised with a Biospec Mini-Beadbeater 8 tissue lyser using four cycles of 30 sec agitation, each followed by a 90 sec cooling period at −20 °C. The homogenised samples were then incubated at −20 °C for 20 min, then heated to 80 °C for 15 min, and returned to −20 °C for at least 20 min. Finally, the samples were centrifuged (21,000 g) at 4 °C for 5 min. The obtained supernatant was filtered (0.45 µm PTFE filters) into internally threaded cryogenic tubes (Fisherbrand) and stored at −80 °C until analysis.

The samples were concentrated using a Savant SC210A SpeedVac Concentrator (ThermoFisher) before derivatisation in a glass vial. Briefly, the samples were reconstituted with 20 µL HPLC (High-performance liquid chromatography)-grade MeOH followed by the addition of 20 µL internal standards, 10 µL 3-nitrophenylhydrazine (3-NPH) at 50 mM in MeOH with 7% pyridine, 10 µL 1-Ethyl-3-(3-dimethylaminopropyl) carbodiimide (EDC) at 50 mM in MeOH with 7% pyridine. The solution was vortexed properly and then incubated for 30 min at 37 °C. In the end, the reaction was quenched by adding 20 µL of water with 0.1% formic acid. The column used in all instances was Kinetex® 2.6 µm XB-C18, 50 × 2.1 mm (Phenomenex, Torrance, CA, USA). The SCFAs profile was quantified using a triple-quadrupole LC-MS/MS (Xevo TQ-S Cronos, Waters), controlled by MassLynx v4.2 software for data acquisition, analysis, and management. All SCFA concentrations (nmol/g) were normalised by tissue weight.

### RNA extraction and cDNA synthesis

Blood samples from 10 fish per sex, dietary regime, and LPS status (negative and positive) were chosen evenly distributed across blocks, resulting in a total of 140 female blood samples and 140 male blood samples (70 female blood samples and 70 male blood samples per LPS status, total n = 280). Total RNA was extracted from each blood sample using the Quick-RNA Whole-Blood kit (Zymo Research), following the manufacturer’s recommendations, with some optimisation to account for the small blood volume. Genomic DNA contamination was removed using the provided DNase I treatment. All RNA concentrations (ng/μL) were quantified using the Qubit RNA HS Assay Kit (Invitrogen, Thermo Fisher Scientific, USA) on a Qubit 4 Fluorometer. In the case of low RNA yield from a blood sample, RNA extraction was performed again from a fresh sample under the same sex, condition, and LPS status. RNA quality was assessed using NanoDrop™ 8000 Spectrophotometer (Thermo Fisher Scientific). All extracted RNA samples were stored at − 80 °C until processing. The amount of RNA from blood samples was normalised to 250 ng per sample. Spleen samples (n = 10 per sex, regime, and LPS) were homogenised by bead beating and total RNA was extracted using the Quick-RNA MiniPrep Plus from Zymo Research. Additionally, the samples were further concentrated using Monarch Spin RNA Cleanup kit (10 µg) (New England BioLabs). The amount of RNA from spleen samples was normalised to 200 ng. cDNA synthesis was performed using the UltraScript 2.0 cDNA Synthesis kit (PCR Biosystems, London, UK) as per the manufacturer’s instructions. All cDNA was diluted 1:5 with molecular-grade water and stored at −80 °C until qPCR.

### Gene expression analysis by qPCR

Primers used in this study are presented in **Table S2**. The primer efficiency of each pair was tested using a seven-point standard curve method^126^. The qPCR experiment was performed using PowerUp™ SYBR™ Green Master Mix for qPCR (Applied Biosystems, US) on the QuantStudio™ 7 Flex Real-Time PCR System (384-well; Applied Biosystems, US). The qPCR cycling conditions were as follows: 1 cycle (50°C for 2 min, then 95°C for 2 min) and 40 cycles (95°C for 15 sec, then 60°C for 1 min). This was followed by the conditions needed for the melt curve analysis as defined by the machine. All primers amplified a single product according to melt curve analysis, and amplicon specificity was confirmed by Sanger sequencing (Source Genomics LTD, Cambridge, CB24 9ZR). All 10 µL qPCR reactions (8 µL master mix* and 2 µL cDNA) were performed in duplicate in white 384-well PCR plates. To minimise any plate effect, all the samples were randomly allocated to the different qPCR plates, keeping an even balance of sex, regime, blocks, and LPS status per plate. To correct for run-to-run plate variations in the blood qPCR experiment, we calculated an inter-run correction for each target gene (*tnfa*, *il1b*, and *il6*) by shifting the C_t_ values to a reference plate. The qPCR results were normalised using the geometric mean of three reference genes: beta-actin (*actb1*), eukaryotic translation elongation factor 1 alpha 1 (*eef1a1*) and ribosomal protein L8 (*rpl8*)^127^. Relative gene expression was first analysed using a linear mixed-effects model, modelling the normalised C_t_ values (ΔC_t_) of the three target genes individually against both fixed and random effects. Then, the comparative (ΔΔCt) method was employed to analyse qPCR data^128^. *(5 µL PowerUp SYBR Green + 0.8 µL 800 nM F. primer + 0.8 µL 800 nM R. primer + 1.4 µL molecular-grade water)

### Genomic DNA extraction, 16S rRNA sequencing, and analysis

From the LPS-negative cohort, intestinal samples from five females and five males were randomly selected from each of the seven regimes, for a total of n = 70 samples. Genomic DNA was extracted using the QIAamp PowerFecal Pro DNA Kit from QIAGEN (Hilden, Germany) as per the manufacturer’s recommendations, which included a bead-beating step using a tissue lyser (Biospec Mini-Beadbeater 8). The extracted genomic DNA was then stored at − 20 °C until processing.

16S rRNA amplicon metagenomic sequencing was performed by Novogene (25 Science Park, Milton, Cambridge, CB4 0FW). Once the quality of the sent genomic DNA samples had been validated, the samples were sequenced according to the Novogene workflow. Briefly, amplicon libraries were produced using universal 16S V3-V4 primers (341F: 5’-CCTAYGGGRBGCASCAG-3’ and 806R: 5’-GGACTACNNGGGTATCTAAT-3’), and sequencing was performed using the paired-end 250 base pairs (bp) Illumina NovaSeq 6000 platform with 50 K tags per sample as data output.

The delivered sequencing data had already been processed by Novogene. After demultiplexing, the amplicon sequence data quality was corrected and denoised using DADA2^129^. Taxonomic annotation was performed using the SILVA 138.1 database^130^. Bioinformatics analysis was conducted using a combination of QIIME2^131^ and phyloseq^132^ in RStudio. α-diversity metrics, namely Observed (Amplicon Sequence Variant, ASV), Chao1, Shannon, and Simpson, were used to assess within-group diversity. Additionally, the effects of “Diet”, “Sex”, and their interaction on unweighted and weighted UniFrac β-diversity were assessed by PERMANOVA (9999 permutations). Differential abundance analysis was performed using ANCOM-BC2^19^.

### Growth metrics and survival parameters

At the beginning and at the end of the 12-week dietary intervention, fish growth metrics and survival parameters were assessed. Briefly, the fish (n = 8) from each experimental tank were transferred to a small breeding tank containing minimal water, which was placed on top of a bench scale (Ohaus, Ranger 3000). The total weight of the fish was recorded after taring the breeding tank and water. Additionally, the number of fish in each experimental tank was recorded at the end of the intervention. Thus, the tanks containing the fish were considered the experimental units and were used to assess growth metrics and survival parameters (**Fig. S12**).

The mean weight per fish in each tank was divided by the total number of fish as follows:

- Initial mean weight per fish in tank = (Initial tank weight)/8 fish per tank
- Final mean weight per fish in tank = (Final tank weight)/ fish survivors

The following formula for the growth metrics was used:

i. Weight difference = (final mean weight) - (initial mean weight)
ii. % weight gain = (weight difference)/ (initial mean weight) X 100
iii. Specific growth rate (SGR) = ([(ln_final mean weight_)-(ln_initial mean weight_)]/ days) X 100

where ln = natural log and days = 84 Survival for each tank was analysed as count data as follows:

- Fish survivors: alive at 12 weeks (between 0 fish and 8 fish)
- Fish dead: 8 - fish survivors

### Statistical analysis

Statistical analyses were performed using R (version 4.6.1)^133^. Unless otherwise stated, descriptive data are presented as mean ± standard error of the mean (SEM). For lmer, degrees of freedom were estimated using the Satterthwaite approximation. Intestinal and brain SCFAs were analysed separately. Concentrations were initially log10-transformed, and linear mixed-effects models were fitted with “Diet”, “Sex”, and their interaction “Diet:Sex” as fixed effects. We applied a Box-Cox transformation to intestinal valerate (λ ≈ −0.34) and to intestinal isovalerate (λ ≈ −0.42), and a Yeo-Johnson transformation to intestinal caproate (λ ≈ −0.10) to meet the assumptions of the linear mixed model. We evaluated model assumptions using appropriate diagnostic procedures. Statistical tests were two-sided, and significance was defined as *p* < 0.05. Where applicable, multiple comparisons were performed and *p* were adjusted for multiple testing. Welch’s two-sample *t*-tests were used to compare tissue weights between sexes. Holm’s method was applied to families of planned contrasts and a Dunnett-type adjustment to comparisons of experimental groups against the Control. Adjusted *p* are reported throughout as *p*_adj_, with multiple comparisons performed as specified.

