## Supplemental Information for "Dietary fibre responses emerge through host-dietary regime interplay: Apple pectin modulates immune reactivity in adult zebrafish"

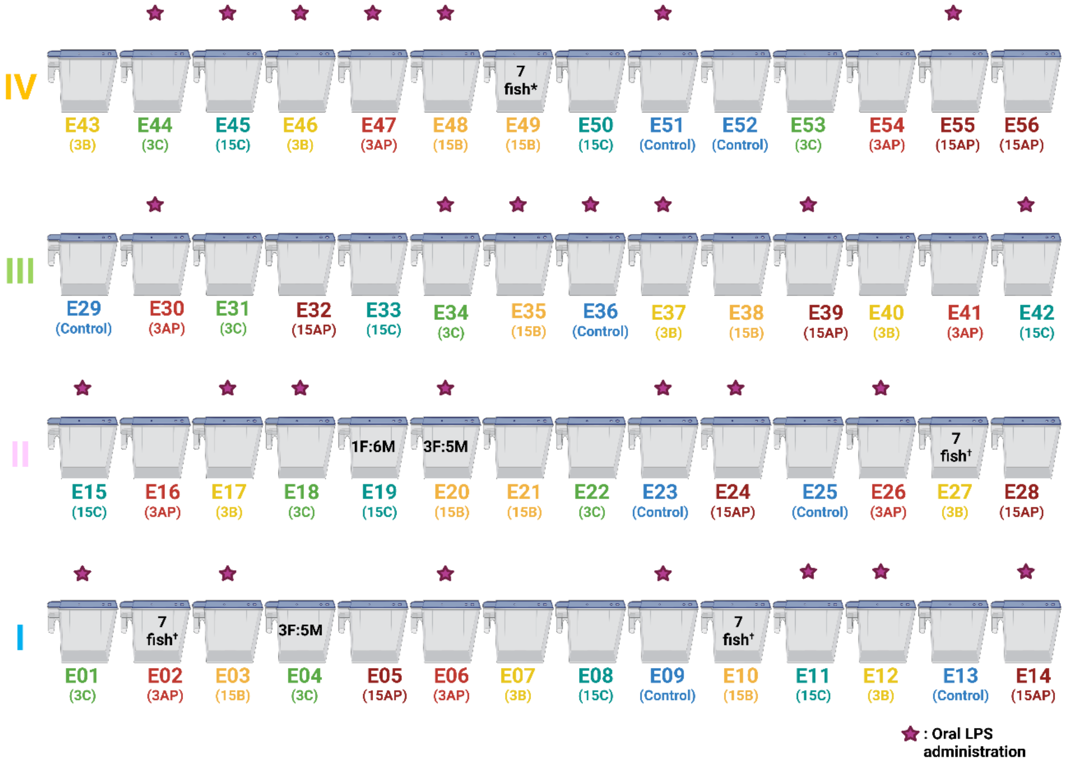


**Fig. S1:** Experimental design set-up.

Female and male zebrafish were housed 1:1 for a total of 8 fish per tank. Roman numerals denote the block number. Each tank was assigned an ID and a dietary regime. †1 dead fish detected at the bottom of the tank. *1 dead fish not found. Created in https://BioRender.com

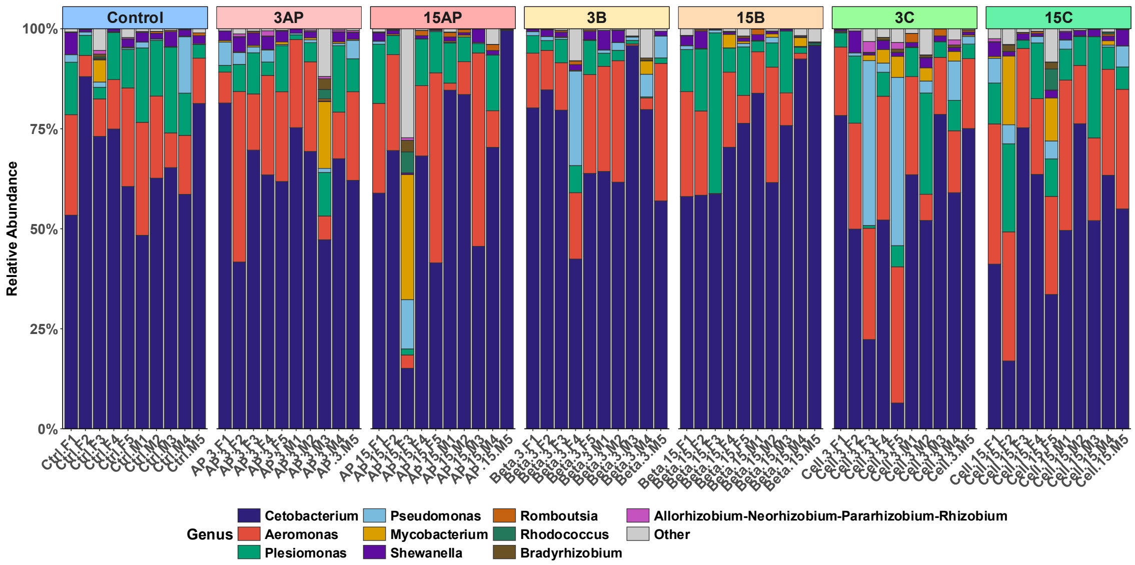


**Fig. S2:** Relative abundance at the genus level.

*Cetobacterium* (phylum: Fusobacteriota) is the most abundant genus (by relative abundance) and a core gut bacterium across all samples.

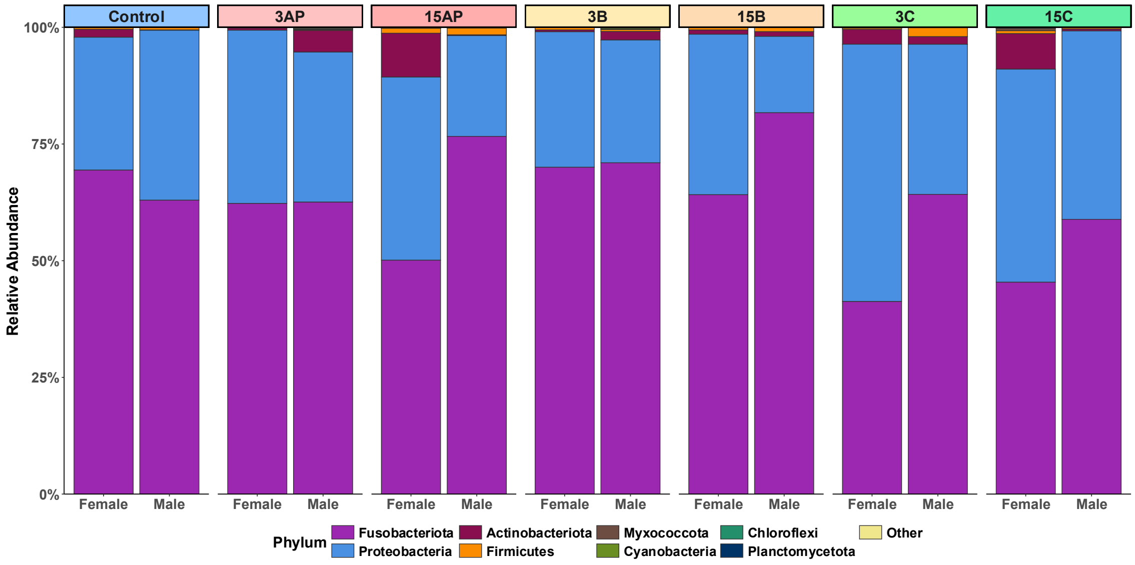


**Fig. S3:** Relative abundance at the phylum level faceted by sex.

Fusobacteriota appear more abundant in males than in females under high-dosage, intermittent dietary regimes.

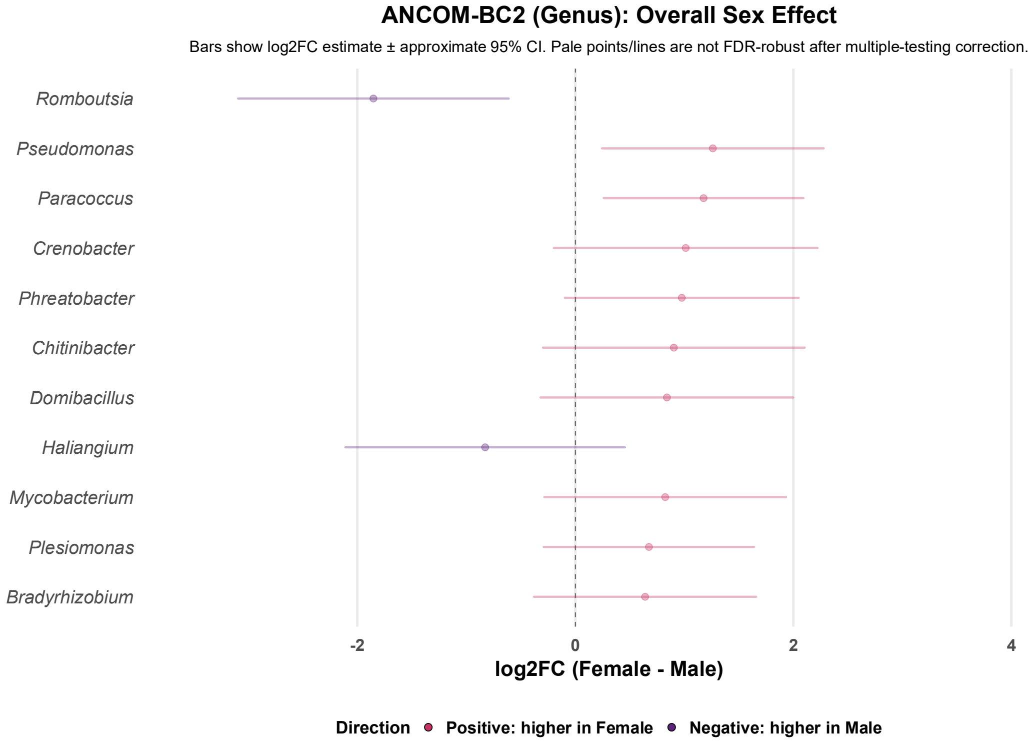


**Fig. S4:** Differential abundance analysis comparing females to males.

Positive values indicate higher estimated abundance in females, whereas negative values indicate higher estimated abundance in males. Genera are ordered by adjusted *p*-value (*q*), with the smallest *q* value at the top; however, no genus-level taxa were differentially abundant after FDR correction ($q > 0.05$).

### Intestine SCFAs — final linear mixed-effects models

Type III F-tests with Satterthwaite denominator degrees of freedom

| Outcome | Term | Num df | Den df | *F* | *p* |
|---|---|---|---|---|---|
| Acetate | Diet | 6 | 21.49 | 2.61 | 0.047 |
| | Sex | 1 | 70.74 | 0.00 | 0.995 |
| | LCMSMS_Run | 1 | 79.43 | 0.00 | 0.945 |
| Butyrate | Diet | 6 | 21.48 | 1.14 | 0.372 |
| | Sex | 1 | 71.32 | 0.48 | 0.493 |
| | LCMSMS_Run | 1 | 2.69 | 0.59 | 0.504 |
| Caproate | Diet | 6 | 22.57 | 0.83 | 0.561 |
| | Sex | 1 | 73.26 | 12.17 | <0.001 |
| | LCMSMS_Run | 1 | 3.15 | 7.87 | 0.064 |
| Isobutyrate | Diet | 6 | 81.71 | 0.48 | 0.823 |
| | Sex | 1 | 82.69 | 10.59 | 0.002 |
| | LCMSMS_Run | 1 | 3.21 | 0.46 | 0.543 |
| Isocaproate | Diet | 6 | 81.88 | 1.93 | 0.085 |
| | Sex | 1 | 82.96 | 9.69 | 0.003 |
| | LCMSMS_Run | 1 | 2.92 | 6.45 | 0.087 |
| Isovalerate | Diet | 6 | 83.00 | 0.77 | 0.595 |
| | Sex | 1 | 83.00 | 0.09 | 0.764 |
| | LCMSMS_Run | 1 | 83.00 | 2.39 | 0.126 |
| 2-Methylbutyrate | Diet | 6 | 83.00 | 0.84 | 0.542 |
| | Sex | 1 | 83.00 | 0.14 | 0.707 |
| | LCMSMS_Run | 1 | 83.00 | 57.78 | <0.001 |
| Propionate | Diet | 6 | 83.00 | 2.65 | 0.021 |
| | Sex | 1 | 83.00 | 0.35 | 0.556 |
| | LCMSMS_Run | 1 | 83.00 | 1.78 | 0.185 |
| Valerate | Diet | 6 | 83.00 | 4.06 | 0.001 |
| | Sex | 1 | 83.00 | 3.87 | 0.052 |
| | LCMSMS_Run | 1 | 83.00 | 4.34 | 0.040 |

**Fig. S5:** Type III ANOVA output for intestinal SCFA linear mixed-effects models.

### Brain SCFAs — final linear mixed-effects models

Type III F-tests with Satterthwaite denominator degrees of freedom

| Outcome | Term | Num df | Den df | *F* | *p* |
|---|---|---|---|---|---|
| Acetate | Diet | 6 | 20.52 | 1.46 | 0.239 |
| | Sex | 1 | 68.16 | 5.41 | 0.023 |
| | LCMSMS_Run | 1 | 1.94 | 1.38 | 0.363 |
| Butyrate | Diet | 6 | 20.96 | 0.72 | 0.640 |
| | Sex | 1 | 70.95 | 2.77 | 0.101 |
| | LCMSMS_Run | 1 | 2.97 | 2.97 | 0.184 |
| Caproate | Diet | 6 | 21.06 | 0.82 | 0.568 |
| | Sex | 1 | 71.26 | 5.15 | 0.026 |
| | LCMSMS_Run | 1 | 3.01 | 1.37 | 0.327 |
| Isobutyrate | Diet | 6 | 21.24 | 0.85 | 0.550 |
| | Sex | 1 | 69.97 | 7.03 | 0.010 |
| | LCMSMS_Run | 1 | 3.26 | 0.78 | 0.438 |
| Isocaproate | Diet | 6 | 20.55 | 0.40 | 0.867 |
| | Sex | 1 | 68.50 | 7.10 | 0.010 |
| | LCMSMS_Run | 1 | 76.48 | 2.41 | 0.125 |
| Isovalerate | Diet | 6 | 19.75 | 0.51 | 0.794 |
| | Sex | 1 | 68.76 | 7.07 | 0.010 |
| | LCMSMS_Run | 1 | 77.78 | 0.31 | 0.581 |
| 2-Methylbutyrate | Diet | 6 | 22.10 | 0.24 | 0.957 |
| | Sex | 1 | 70.29 | 5.60 | 0.021 |
| | LCMSMS_Run | 1 | 3.22 | 9.07 | 0.052 |
| Propionate | Diet | 6 | 21.05 | 0.94 | 0.485 |
| | Sex | 1 | 68.49 | 6.27 | 0.015 |
| | LCMSMS_Run | 1 | 75.92 | 2.90 | 0.093 |
| Valerate | Diet | 6 | 21.41 | 0.44 | 0.845 |
| | Sex | 1 | 69.44 | 2.32 | 0.133 |
| | LCMSMS_Run | 1 | 77.33 | 0.04 | 0.833 |

**Fig. S6:** Type III ANOVA output for brain SCFA linear mixed-effects models.

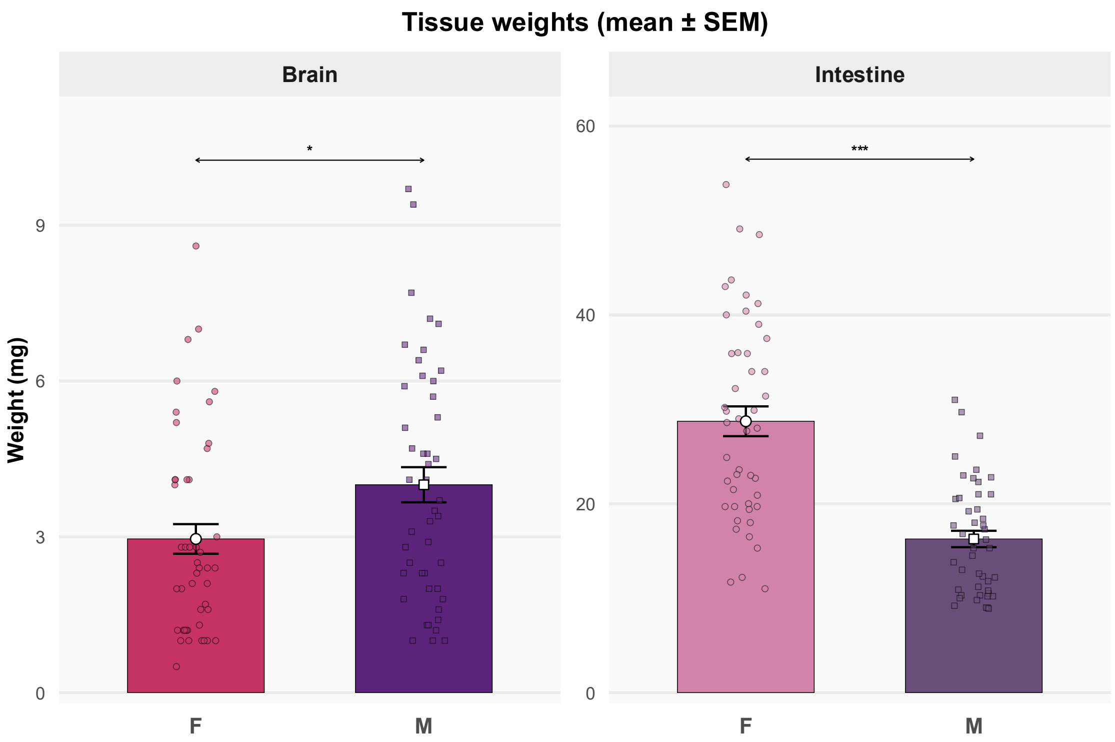


**Fig. S7:** Brain and intestine mean weights by sex.

Bar charts displaying the mean weights (mg) of the brain and intestinal samples in our study, grouped by sex. The centre dots and error bars indicate the mean ± standard error of the mean. Each individual dot represents a unique sample. F denotes females, and M denotes males. *denotes $p = 0.02$ and ***denotes $p < 0.001$.

| Diet | Sex | Tissue | n |
|---|---|---|---|
| Control | F | Intestine | 7 |
| Control | F | Brain | 7 |
| Control | M | Intestine | 6 |
| Control | M | Brain | 6 |
| 3AP | F | Intestine | 7 |
| 3AP | F | Brain | 7 |
| 3AP | M | Intestine | 6 |
| 3AP | M | Brain | 6 |
| 15AP | F | Intestine | 7 |
| 15AP | F | Brain | 7 |
| 15AP | M | Intestine | 7 |
| 15AP | M | Brain | 7 |
| 3B | F | Intestine | 7 |
| 3B | F | Brain | 7 |
| 3B | M | Intestine | 7 |
| 3B | M | Brain | 7 |
| 15B | F | Intestine | 6 |
| 15B | F | Brain | 6 |
| 15B | M | Intestine | 7 |
| 15B | M | Brain | 7 |
| 3C | F | Intestine | 7 |
| 3C | F | Brain | 7 |
| 3C | M | Intestine | 7 |
| 3C | M | Brain | 7 |
| 15C | F | Intestine | 5 |
| 15C | F | Brain | 5 |
| 15C | M | Intestine | 6 |
| 15C | M | Brain | 6 |

**Fig. S8:** Metabolomics sample sizes by diet, sex, and tissue

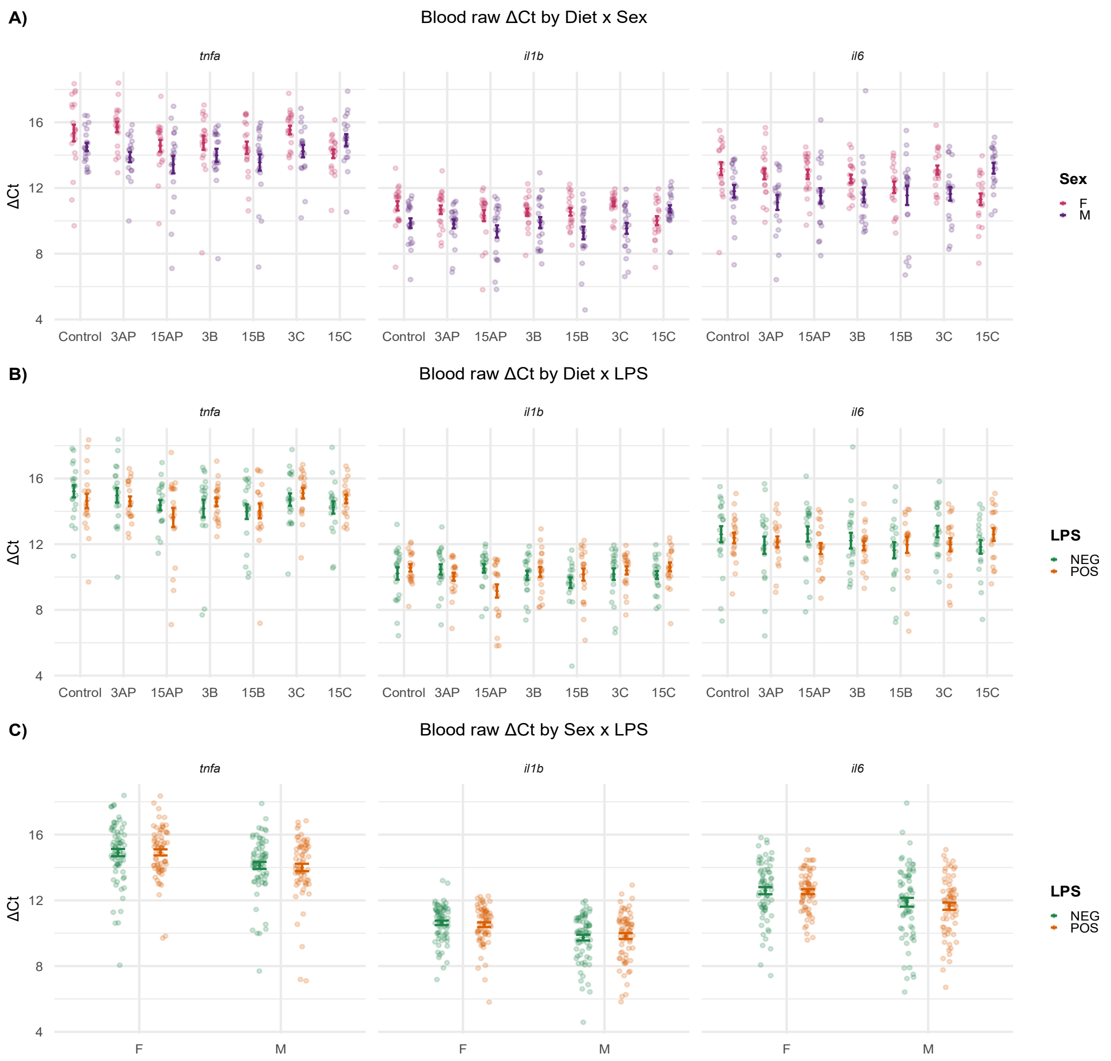


**Fig. S9:** Normalised blood gene expression of target genes.

Blood raw ΔCt values of the target genes (*tnfa*, *il1b*, and *il6*) normalised to the three reference genes (*actb1*, *eef1a1*, and *rpl8)*. The centre dots and error bars indicate the mean ± standard error of the mean. F denotes females, and M denotes males. (A) The plot shows the interactions between "Diet" and "Sex". (B) The plot shows the interactions between "Diet" and "LPS". (C) The plot shows the interactions between "Sex" and "LPS".

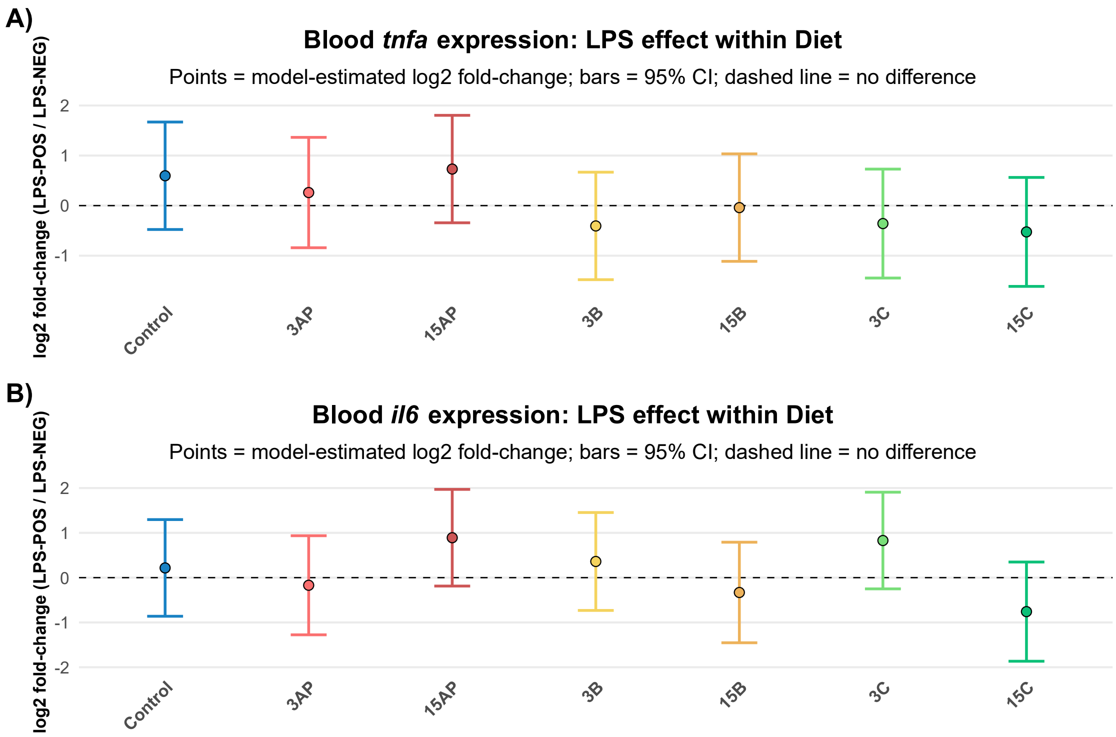


**Fig. S10:** Blood *tnfa* and *il6* expression testing the effects of LPS.

**(A-B)** Gene expression patterns of *tnfa* and *il6*, comparing LPS POS vs LPS NEG under each of the seven dietary regimes. Each point is the model-estimated log2FC, and the vertical error bar is its 95% confidence interval.

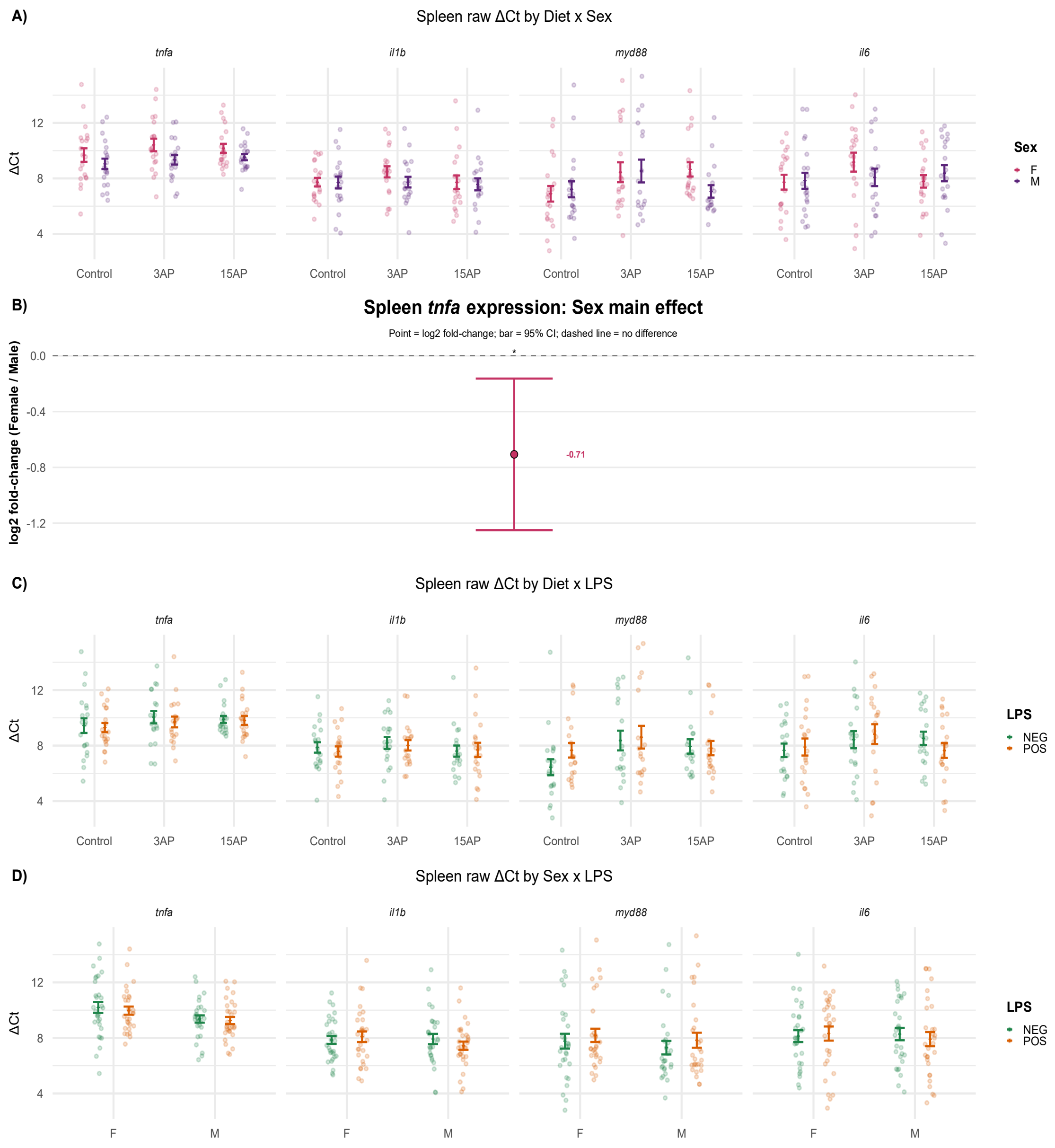


**Fig. S11:** Normalised spleen gene expression of target genes.

**(A, C, D)** Spleen raw ΔCt values of the target genes (*tnfa*, *il1b*, *myd88*, and *il6*) normalised to the three reference genes (*actb1*, *eef1a1*, and *rpl8)*. The centre dots and error bars indicate the mean ± standard error of the mean. F denotes females, and M denotes males. **(A)** The plot shows the interactions between "Diet" and "Sex". **(B)** Female spleen *tnfa* expression lower than males. **(C)** The plot shows the interactions between "Diet" and "LPS". **(D)** The plot shows the interactions between "Sex" and "LPS". *denotes $p = 0.01$.

| Diet summary (mean ± SEM; tank = experimental unit) | | | | | | | |
|---|---|---|---|---|---|---|---|
| Diet | n | Initial mean weight (g) | Final mean weight (g) | Weight difference (g) | Percentage gain (%) | SGR (%/day) | Survival (%) |
| Control | 8 | 0.510 ± 0.025 | 0.683 ± 0.033 | 0.173 ± 0.039 | 35.96 ± 9.17 | 0.348 ± 0.076 | 100.00 ± 0.00 |
| 3AP | 8 | 0.535 ± 0.043 | 0.678 ± 0.039 | 0.143 ± 0.051 | 31.49 ± 12.06 | 0.293 ± 0.104 | 98.44 ± 1.56 |
| 15AP | 8 | 0.536 ± 0.021 | 0.641 ± 0.024 | 0.105 ± 0.025 | 20.38 ± 5.05 | 0.213 ± 0.051 | 100.00 ± 0.00 |
| 3B | 8 | 0.497 ± 0.031 | 0.711 ± 0.028 | 0.214 ± 0.039 | 46.58 ± 10.13 | 0.435 ± 0.082 | 98.44 ± 1.56 |
| 15B | 8 | 0.503 ± 0.031 | 0.650 ± 0.028 | 0.147 ± 0.040 | 32.18 ± 9.09 | 0.313 ± 0.081 | 96.88 ± 2.05 |
| 3C | 8 | 0.491 ± 0.024 | 0.681 ± 0.039 | 0.190 ± 0.028 | 39.09 ± 6.01 | 0.385 ± 0.053 | 100.00 ± 0.00 |
| 15C | 8 | 0.507 ± 0.024 | 0.668 ± 0.031 | 0.162 ± 0.042 | 34.40 ± 10.47 | 0.330 ± 0.083 | 100.00 ± 0.00 |

**Fig. S12:** Growth parameters and survival percentage.

The main effect of "Diet" was not statistically significant on the weight difference ($F_{6,46} = 1.1$, $p = 0.37$), % weight gain ($F_{6,46} = 0.94$, $p = 0.48$), nor the SGR ($F_{6,46} = 1.01$, $p = 0.43$). Moreover, neither of the three calculated growth metrics were significantly different when comparing each experiment diet to Control ($p_{adj} > 0.05$). Additionally, survival was uniform across all the seven regimes (Bias-reduced logistic regression: Diet: $\chi^2 = 5.93$, df = 6, $p = 0.43$).

**Table S1:** Calculated approximate composition of diets.

| | **Control basal*** | | **3DF** | **15DF** |
|---|---|---|---|---|
| **Composition** | Fish Meal, Vital wheat gluten, Fish protein hydrolysed, Crustacea meal, Pea protein, Gelatine, Algal oil (*Schizochytrium* sp.), Fish oil. | | Fish Meal, Vital wheat gluten, Fish protein hydrolysed, Crustacea meal, Pea protein, Gelatine, Algal oil (*Schizochytrium* sp.), Fish oil, 3% DF[!]. | Fish Meal, Vital wheat gluten, Fish protein hydrolysed, Crustacea meal, Pea protein, Gelatine, Algal oil (*Schizochytrium* sp.), Fish oil, 15% DF[!]. |
| **Diet** | **Final DF inclusion**** | **Protein g/100 g final diet** | **Fat g/100 g final diet** | **Total carbohydrate (including DF) g/100 g final diet** |
| Control | 0% | 62.0 | 13.0 | 7.0 |
| 3AP | 2.9% | 60.2 | 12.6 | 9.2-9.4 |
| 3B | 2.9% | 60.2 | 12.6 | 9.6 |
| 3C | 2.9% | 60.2 | 12.6 | 9.6-9.7 |
| 15AP | 13% | 53.9 | 11.3 | 16.9-17.8 |
| 15B | 13% | 53.9 | 11.3 | 18.6 |
| 15C | 13% | 53.9 | 11.3 | 18.5-19.1 |

*Nutritional additives: Vitamins, pro-vitamins and chemically well-defined substances having a similar effect (per kg): vitamin D3 (3a671) 3000 IU; vitamin E (3a700) 1.2 g; vitamin C (3a311) 1.9 g; betaine (3a925) 10.8 g. Compounds of trace elements (mg/kg): iron (3b103) 9; iodine (3b201) 0.8; zinc (3b605) 11.3; copper (3b405) 13.5; manganese (Eb502) 14.4. The exact percentage composition of ingredients is not disclosed by Sparos.

** 3 g DF + 100 g basal = 2.9% w/w final diet and 15 g DF + 100 g basal = 13.0% w/w final diet

[1]DF: The dietary fibres used were purchased as analytical-grade reagents rather than food-grade ingredients. Approximate composition values were estimated from supplier specifications/literature and expressed as g/100 g ingredient as supplied. The carbohydrate fraction was assumed to represent predominantly dietary fibre polysaccharides.

Apple pectin (Sigma): ~ 0 g protein, ~ 0 g fat, ~ 83-90 g total carbohydrate

Oat β-glucan (Megazyme): ~ 0.1 g protein, ~ 0 g fat, ~ 95.8 g total carbohydrate

Cellulose (Sigma): ~ 0 g protein, ~ 0 g fat, and ~ 95-100 g total carbohydrate

**Table S2:** qPCR primer pairs used in this study.

| **Gene** | **Primer sequence (5′-3′)** | **Melting Temp (Tm °C)** | **Efficiency (%)** | **Amplicon length** | **Accession number** | **Ref** |
|---|---|---|---|---|---|---|
| *actb1* | F: ATGGATGAGGAAATCGCTG<br>R: ATGCCAACCATCACTCCCTG | F: 62.7<br>R: 67.2 | 114 | 108 bp | BC045846.1 | [1] |
| *eef1a1a* | F: ATCGACCGTCGTTCTGGCAAG<br>R: AACGGTGTGATTTGAGGGAA | F: 70.6<br>R: 64.3 | 104.5 | 250 bp | AY422992.1 | [2] |
| *rpl13a* | F: TGGTGAGGTGTGAGGGTATCAAC<br>R: AATTTGCGTGTGGGTTTCAGAC | F: 67.1<br>R: 67 | 99 | 269 bp | NM_212784.1 | [2] |
| *tnfa* | F: GCTGGATCTTCAAAGTCGGGTGTA<br>R: TGTGAGTCTCAGCACACTTCCATC | F: 69.1<br>R: 67.8 | 104 | 107 bp | AY427649.1 | [1] |
| *il1b* | F: CTCAGCCTGTGTGTTTGGGA<br>R: GGGACATTTGACGGACTCG | F: 66.4<br>R: 65.6 | 108.6 | 176 bp | NM_212844.2 | [3] |
| *il6* | F: AGACCGCTGCCTGTCTAAAA<br>R: TTTGATGTCGTTCACCAGGA | F: 63.8<br>R: 64.2 | 106.5 | 107 bp | NM_001261449.1 | [1] |
| *myd88* | F: GATGATCCACAGGGACTGACACC<br>R: GGAAGGACGTCTCTGTCAAACAC | F: 69<br>R: 66 | 107.75 | 129 bp | NM_212814.2 | [4] |

The qPCR primer pairs were obtained from the literature. The melting points were obtained from the supplier (Sigma). The primer efficiencies were obtained from a seven-point dilution standard curve. The accession numbers were obtained by BLAST (Basic Local Alignment Search Tool). The amplicon lengths were obtained from Sanger sequencing (Source Genomics LTD, Cambridge).
